# Duration judgments with conflicting audiovisual cues

**DOI:** 10.64898/2026.08.19.745628

**Authors:** Ömer F. Yildiran, Long Ni, Michael S. Landy

## Abstract

Previous work showed that observers integrate audiovisual duration cues optimally when cue-conflict is small. Does causal inference lead to a breakdown of audiovisual integration when duration conflicts are large? We addressed this by testing a wide range of duration cue-conflicts. Participants compared the auditory durations of a test and a standard stimulus. Audiovisual durations were consistent in the test stimulus, but differed by seven conflict durations (up to 250 ms) in the standard. Two levels of auditory noise were tested. Auditory duration percepts shifted systematically toward the visual duration, especially with high auditory noise. The shift was proportional to cue-conflict magnitude, inconsistent with causal inference. We compared several models. A heuristic model in which the observer probabilistically switches between the visual and auditory cues was preferred for most participants, although performance differences across models were small. Within the tested conflict range, the forced fusion, causal inference, and probabilistic cue switching models produced overlapping, near-linear shifts as a function of cue-conflict. Model simulations further revealed that given the measured sensory noise, forced fusion and causal inference can be discriminated only with unreasonably large conflicts. Together, while our results suggest that observers do not rely on causal inference when judging auditory durations under our conditions, high sensory encoding noise in auditory duration limits the discriminability of competing computational models.

## Introduction

Accurately estimating the timing of events is essential for many everyday behaviors, from playing a musical instrument to catching a fly ball. Humans rely on multiple sensory modalities to perceive the timing and duration of events in the world.

Prevailing models of timing mechanisms often rely on a clock analogy, particularly pacemaker– accumulator models (Treisman, 1963; Gibbon, 1977; Wittmann, 2009; Grondin, 2014; Hartcher-O’Brien et al., 2016; Grondin, 2025). These models propose that the brain contains an internal clock, where a pacemaker emits regular pulses that are accumulated over time to estimate duration. However, there is no concrete evidence for the existence of such a mechanism in the brain. While these models successfully explain timing properties such as Vierordt’s law (Lejeune and Wearden, 2009; Jazayeri and Shadlen, 2010; Mamassian and Landy, 2010), which accounts for overestimation of short durations and underestimation of long durations, neural response patterns predicted by these models do not align well with actual neural data (Buonomano and Laje, 2010; Muller and Nobre, 2014; Kononowicz and van Wassenhove, 2016). Unlike vision or audition, however, time does not have a dedicated sensory receptor; rather, temporal perception is inferred based on signals elicited from multiple modalities.

Despite the inherent difficulty of temporal estimation, the brain must operate with remarkable precision and consistency when timing our actions, which is crucial for effective behavior. Consider ensemble music performance: after hearing a musical phrase, you must time your entrance so it coincides with the other performers, and entering too early or too late can disrupt the coordination of the group. Achieving this kind of temporal precision relies not only on motor coordination but also on the brain’s ability to accurately perceive and integrate temporal cues from multiple senses.

In natural environments, sounds and sights rarely occur in isolation; temporal judgments should often be made across modalities. Continuing our musical example, estimation of when you should begin to play can come both from hearing the violinist and from seeing the movements of her bow. This multisensory encoding makes time perception inherently more variable and harder to investigate. The brain estimates time using cues embedded in visual, auditory, and possibly from other sensory inputs, each with its own level of precision and bias. Even when stimuli are relatively clear (low external noise), differences in the physical properties and neural conduction velocities of vision and audition create inherent temporal discrepancies. Light travels faster than sound, but paradoxically, auditory signals often reach the cortex more quickly than visual ones due to faster transduction and shorter conduction delays (Zoefel and VanRullen, 2017). Auditory stimuli typically arrive at central processing regions within 8–10 ms, whereas visual signals take approximately 20–40 ms to reach the cortex from the retina (King and Palmer, 1985; Vroomen and Keetels, 2010). In this study, we investigated how the brain perceives durations when presented with conflicting information across modalities, specifically examining whether and how observers integrate auditory and visual timing cues under such conflict.

When sensory information is available from multiple modalities, the brain faces the challenge of integrating these cues into a single percept. Research across diverse sensory systems has shown that this integration is not arbitrary. Rather, the brain tends to weight each cue according to its reliability. This strategy—known as statistically optimal cue integration—helps minimize the total uncertainty of the fused percept (Maloney and Landy, 1989; Young et al., 1993; Landy et al., 1995; Ernst and Banks, 2002). This optimal cue-integration strategy has been documented across a wide array of sensory tasks, including vision-touch integration for object slant estimation (Ernst and Banks, 2002; Hillis et al., 2002), taste-smell combination for flavor perception (Maier and Elliott, 2020), and audiovisual integration for spatial localization (Alais and Burr, 2004; Negen et al., 2018). These findings underscore the generality of Bayesian cue combination as a core principle of sensory processing (Landy and Kojima, 2001; Landy et al., 2011; Wolpert et al., 2011; Alais and Burr, 2019; Rossi et al., 2021), providing a robust normative framework for understanding multisensory perception.

In previous experiments (Shams et al., 2005; Hartcher-O’Brien et al., 2014), sensory cues across modalities were deliberately aligned in space and time, or only slightly mismatched, effectively biasing the brain toward assuming a single source. The use of low-conflict conditions is a way of examining how different cues are combined with respect to their reliabilities. Under such low-conflict conditions, it is reasonable to assume a common cause and therefore integrate the cues. However, everyday perception often presents greater cue disparities, whether from environmental factors (e.g., room acoustics), technological delays (e.g., audio–video lag), or neural processing differences. In those cases, the brain must address two interrelated questions: Are these disparate cues from the same source? And if so, how should they be combined? Rather than integrating across the board, the brain may dynamically shift from fusion to segregation depending on the degree of mismatch, an insight that has prompted the development of Bayesian causal-inference models. These models explain how integration diminishes as spatial or temporal disparity grows, as notably demonstrated in the ventriloquist-illusion paradigm (Shams et al., 2005; Körding et al., 2007; Hong et al., 2021). Under this account, integration is conditional on inferred causal structure rather than obligatorily applied to all instances of cue combination (Körding et al., 2007; Badde et al., 2018, 2021; Hong et al., 2021; Badde et al., 2023; Li et al., 2025).

While causal inference has been implicated in multisensory perception across various sensorimotor tasks, much less is known about whether it operates in temporal perception, particularly in duration judgments. Bayesian causal-inference models posit that perception emerges from a combination of common-cause inference and likelihood-based integration; that is, the brain infers whether different sensory cues share a source and, if so, integrates them accordingly. Hartcher-O’Brien et al. (2014) provided evidence for optimal cue integration in duration perception under low-conflict conditions: when audiovisual signals specified the similar durations, observers combined duration information in a statistically optimal fashion. However, it remains unexplored whether visual and auditory duration are still integrated when the conflict between them becomes substantial, as Bayesian causal inference predicts a breakdown of cue integration under large-conflict conditions.

In the present study, we investigated how observers judge auditory duration when presented with conflicting audiovisual (AV) cues. We manipulated two key factors: the degree of cross-modal cue conflict and auditory cue reliability. Before introducing audiovisual cue conflict, we measured each participant’s audio-visual temporal bias in a cross-modal duration-discrimination task and compensated for it in the bimodal experiment. Successful manipulation of auditory cue reliability was confirmed in a separate unimodal experiment. Although participants were instructed to base their judgments on the auditory signal alone, perceived auditory duration was systematically biased toward the task-irrelevant visual duration, especially when the auditory cue was less reliable. Moreover, we observed that visually induced biases in auditory duration judgments increased approximately linearly with audiovisual conflict magnitude. This pattern differs from the non-linear breakdown of integration predicted by traditional Bayesian causal-inference models. Instead, the observed behavior was consistent with a model in which observers flexibly switched between reliance on auditory and visual cues across trials. Model comparison further suggested that, under substantial sensory uncertainty, multiple computational strategies produce highly similar behavioral predictions, limiting the discriminability of competing integration models.

With well-controlled experiments and a systematic evaluation of computational models, this work characterizes auditory duration perception under substantial audiovisual cue conflict and highlights the challenges in studying multisensory time perception.

## Methods

### Participants

We recruited thirteen participants for the experiment, two of whom were the authors O.Y. and L.N., two were other lab members, while the rest were naive to the purposes of the experiment but experienced in psychophysical experiments. The study was conducted in accordance with the guidelines laid down in the Declaration of Helsinki and approved by the New York University Institutional Review Board (reference number: IRB-2016-595). All participants provided informed consent prior to their involvement in the study and were compensated $15 per hour for participation, except for the first author.

### Apparatus

All experiments were conducted on a 21-inch CRT (Trinitron GDM-5402, Sony, Japan) with a refresh rate of 120 Hz and a spatial resolution of 800×600 pixels. The experiments were designed and executed using the Python PsychoPy package (Peirce, 2007). The participants sat in a darkened room, approximately 60 cm away from the computer screen, with their head stabilized by a chin rest.

### Stimuli

#### Visual stimuli

The visual stimulus was a black disk (0.07 cd/m^2^) displayed in the center of the screen on a gray background (8.8 cd/m^2^). The width of the disk was 3 degrees of visual angle. Throughout each trial the disk’s black outline was continuously displayed. During the stimulus intervals, the inside of the circle changed from the background gray to black.

#### Auditory stimuli

We adapted the stimuli from Hartcher and colleagues (2014) and Shi and colleagues (2013) in which they created their stimuli by adding a pure-tone signal to a pink-noise background and manipulated duration uncertainty by varying the signal-to-noise ratio. Our stimulus consisted of the sum of a bandpass *background* noise that had a constant level throughout the trial and a second bandpass *signals* noise (Figure 1A). The signals waveform had a low, non-zero amplitude before, between and after the pair of standard and test intervals. The duration of the standard interval was fixed at 500 ms and the duration of the test interval varied over trials. During the standard and test intervals, the signal amplitude was increased (5 ms cosine ramp up, flat peak, cosine ramp down). We differentiated the signal from the background using different broadband filters to ensure that the test and standard intervals were always detectable even when the background noise was louder than the signal. The background was band-pass filtered with cutoffs at 10 and 610 Hz. The signal waveform (including the test and standard) was band-pass filtered from 150–775 Hz, both sampled at 48 kHz. All filtering was implemented using second-order sections (SOS) Butterworth filters with a filter order of 4.

**Figure 1.**
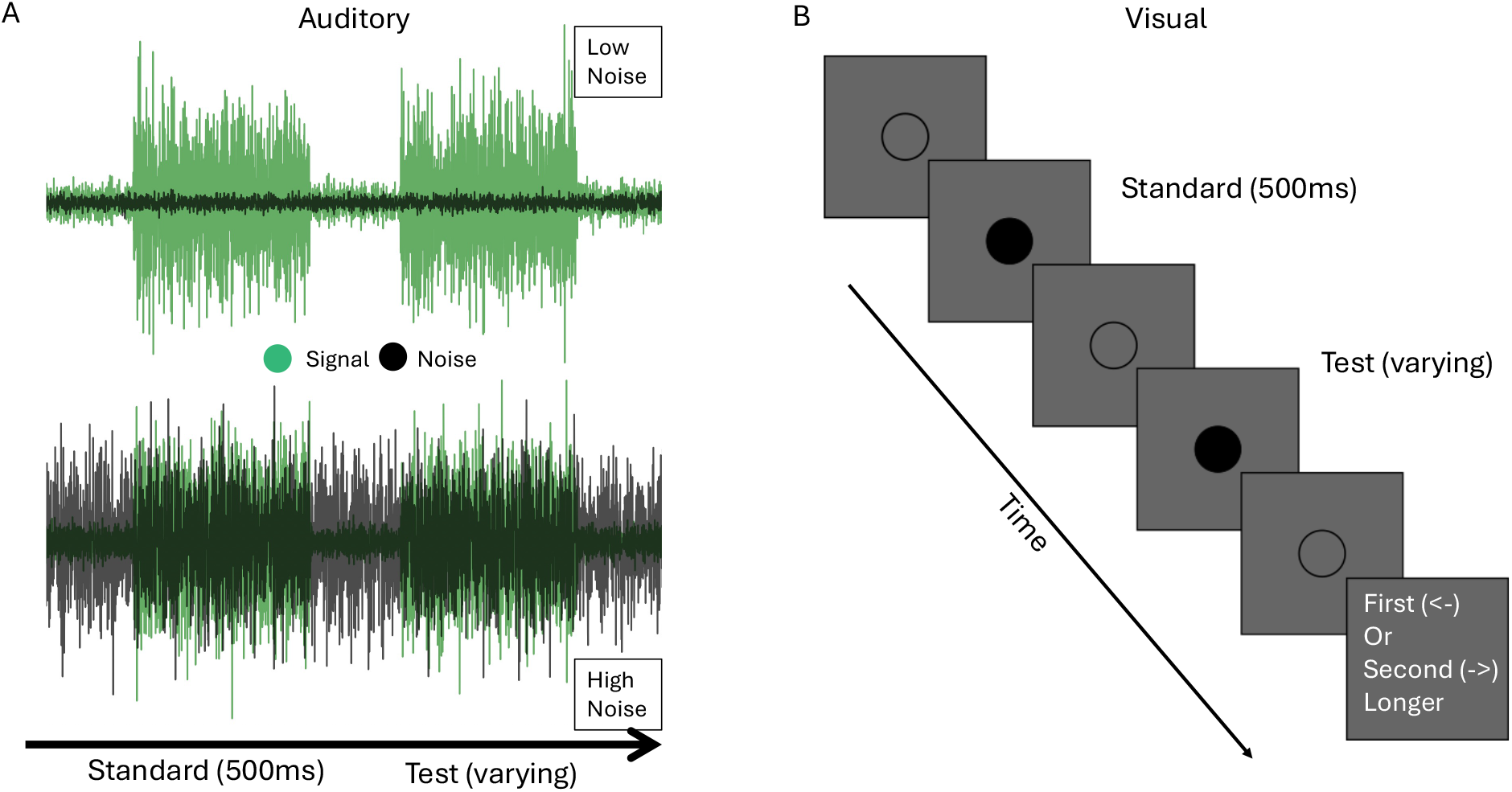
Experimental stimuli and trial procedure. Schematic illustration of the auditory and visual stimuli used in the duration-discrimination tasks. (**A**) Example trials of the unimodal auditory task under the low (top) and high (bottom) auditory-noise conditions. In the low-noise condition, the auditory signal (bright green) is clearly distinguishable from the background noise (black). In the high-noise condition, signal and noise overlap substantially. (**B**) Trial timeline for the unimodal visual task. A disk outline remained visible throughout the trial to provide a stable visual reference. Each trial consisted of two sequentially presented stimulus durations (a standard and a test), separated by a short inter-stimulus interval (ISI). During each signal duration, the disk was filled (signal-on; shown dark), indicating the presence of the visual stimulus. Pre- and post-stimulus periods flanked the two intervals. The 500-ms standard and an adaptively varied test duration were presented in random order (standard-first order shown). Participants reported whether the first or second duration was longer.

We manipulated auditory uncertainty by varying the amplitude of the background relative to the signals waveform before finally summing the two waveforms. In the low-noise condition, the background amplitude was approximately 0.1 times the signal amplitude, whereas in the high-noise condition it was approximately 1.2 times that amplitude. The final mixed waveform was then peak-normalized for playback.

Auditory stimuli were calibrated using a Bruel & Kjaer Type 2240 sound level meter positioned at the participant’s head location. Sound levels were measured as A-weighted equivalent continuous levels (LAeq). During the test and standard intervals the summed waveforms had an average intensity of approximately 70.5 dB SPL. In the low-noise condition, during the non-stimulus intervals the sum of the background waveform and the non-stimulus waveform had an amplitude of 57.4 dB SPL, yielding a signal-to-noise ratio (SNR) of approximately +13.1 dB. In the high-noise condition, the non-stimulus waveform had an amplitude of 67.5 dB SPL, yielding an SNR of approximately +2.9 dB. All sound intensities are reported as A-weighted levels.

To further characterize the stimulus structure, we additionally measured the sound levels of the individual stimulus components prior to mixing the background and signal waveforms. The background waveform averaged approximately 51.5 dB SPL in the low-noise condition and 67.0 dB SPL in the high-noise condition. For the signal waveform, the level during non-stimulus intervals (pre-stimulus, ISI, post-stimulus) was approximately 57.1 dB SPL in the low-noise condition and 54.5 dB SPL in the high-noise condition. During the stimulus intervals, levels increased to approximately 70.6 dB SPL in the low-noise condition and 66.6 dB SPL in the high-noise condition. These measurements confirm that the stimulus contained a clearly detectable signal embedded in background noise, with the signal interval producing a distinct increase in sound level relative to the baseline.

### Procedure

Participants completed four experiments. First, they completed unimodal auditory and unimodal visual tasks, which were used to measure the uncertainty of duration perception for each modality. They carried out these two tasks in separate blocks at the beginning of the experiment to become familiar with the stimuli.

After completing the unimodal tasks, participants completed the cross-modal baseline task, allowing us to measure and correct for the perceptual bias between auditory and visual stimulus durations. One participant did not complete the main experiment, as their cross-modal bias was too large to be reliably calibrated (a 560 ms bias toward visual duration in the high-noise auditory condition), and a second participant was excluded due to a procedural difference in an early version of the experiment. All subsequent analyses were therefore conducted on the remaining eleven participants.

Finally, participants performed the main experiment, which consisted of three sessions of approximately one hour each. Each session started with a flexible number of practice trials in which participants received feedback about the correctness of their answers until they felt comfortable with each task.

All tasks used a 2IFC design (“Which interval was longer?”). In each trial there was a standard and a test stimulus. The order of standard/test stimuli was randomized across trials. Each trial comprised a prestimulus noise, first stimulus, an inter-stimulus interval (ISI), second stimulus, and a post-stimulus noise. Short breaks occurred every 30 trials.

In the unimodal auditory, unimodal visual, and cross-modal experiments, the standard duration was fixed at 500 ms. In the bimodal experiment, the auditory component of the standard interval was always 500 ms. When audiovisual conflict was 0 ms, the visual component was also nominally 500 ms, but was adjusted based on the participant’s audiovisual duration bias (see below). For non-zero conflict conditions, the visual duration was adjusted on each trial according to the conflict level (e.g., +250 ms conflict corresponded to a 750 ms nominal visual duration), while the auditory duration remained fixed at 500 ms.

Test stimulus duration was varied across trials according to adaptive staircase procedures (1-up/3-down or 1-down/3-up in the bimodal experiment). In the bimodal experiment, we compensated for each participant’s modality-specific duration bias, measured in the cross-modal task, by adjusting the visual stimulus duration. This subject-specific correction ensured that auditory and visual components were perceptually matched in the test stimuli and in the zero-conflict standard condition. Below, we describe each experiment in detail.

The duration of the visual stimulus was adjusted to perceptually match the auditory duration in both intervals. Then the conflict duration was added to the visual duration in the standard interval. Each trial consisted of a pre-stimulus interval (200–450 ms), the first stimulus (standard or test), an ISI (400-900 ms), the second stimulus (test or standard), and a post-stimulus interval (200-450 ms), forming one continuous audiovisual stimulus.

#### Unimodal baselines (Auditory only, Visual Only)

In the unimodal tasks, 11 participants performed a 2IFC duration-discrimination task, in which either two auditory durations (i.e., signals embedded on a background noise) or two visual durations (i.e., visual filled disks as signals) were sequentially presented. The auditory task lasted approximately 20 minutes, and the visual task lasted around 12 minutes.

In both tasks, the standard stimulus duration was fixed at 500 ms. The duration of the test auditory stimulus varied (between 25 and 975 ms) from trial to trial. Waveforms of example stimuli in the low and high auditory-noise conditions are shown in Figure 1.

Test duration was determined using four interleaved staircases: 1-Down 2-Up, 1-Up 2-Down, 1-Down 3-Up, and 1-Up 3-Down staircases of 70 trials each, resulting in a total of 280 trials for each unimodal task. We also included 14 trials with a large duration difference between the standard and test durations (+0.45 *s* or −0.45 *s*) for each task. These trials allowed us to better estimate the lapse rate (Prins, 2012).

#### Cross-modal duration comparison

We ran a cross-modal duration-comparison task in order to estimate any participant-specific bias in the perception of visual vs. auditory temporal intervals (Figure 2). In this task, participants compared the durations of one auditory and one visual stimulus. In each trial, the presentation order was pre-cued. We split the experiment into 30 mini blocks of 10 trials and the last mini block of 8 trials (a total of 308 trials). These trials corresponded to two 70-trial staircases (one 1-up 1-down and one 1-down 1-up) for each of the two conditions (low and high auditory noise), and 14 easy lapse-rate trials for each condition. The visual stimulus (standard) was fixed at 500 ms while the auditory stimulus duration varied based on the staircase. The noise conditions were intermixed. Before each mini block, we informed participants about the order of stimuli (e.g., “In this block the order of modalities will be: V->A”) to orient attention to both intervals and to help them understand the nature of the task. The order of stimulus presentation was randomized across mini-blocks.

**Figure 2.**
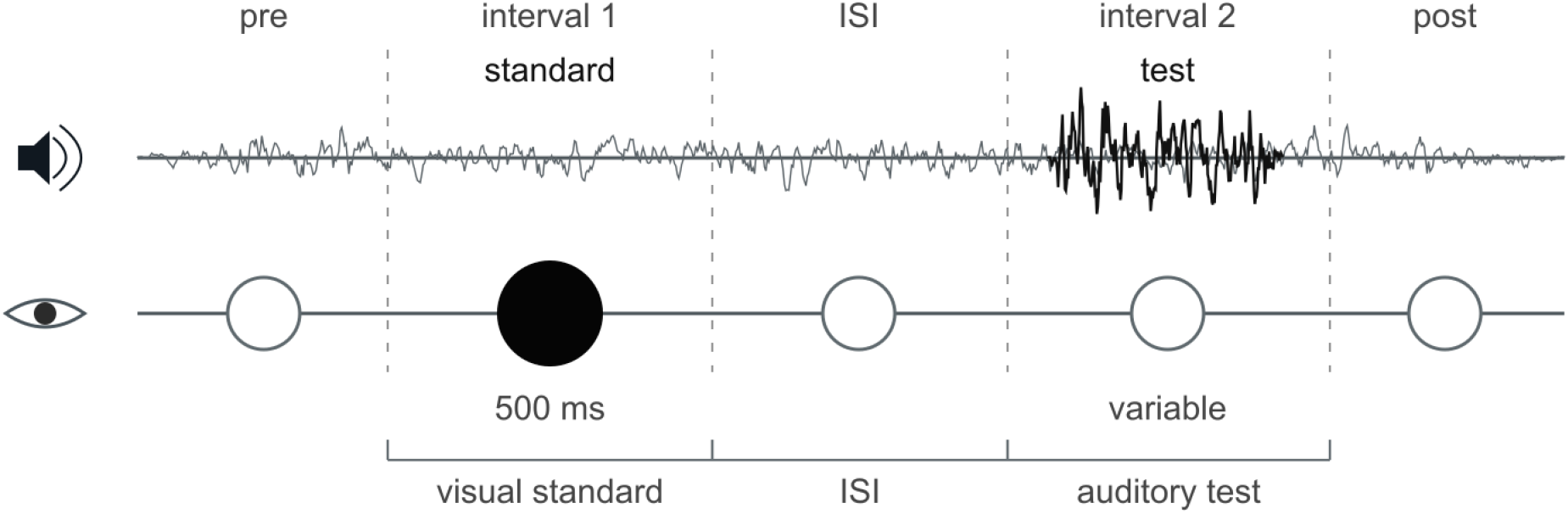
Cross-modal duration-comparison trial structure. Schematic of a single cross-modal trial. A trial consisted of two sequential stimuli (standard and test), separated by an inter-stimulus interval (ISI, dashed vertical lines indicate interval boundaries). **Top (auditory):** Continuous background noise was presented throughout the trial. A signal burst occurred during either the first or second interval. **Bottom (visual):** A disk outline was continuously visible. During the signal duration, the disk was filled. In this task, the visual stimulus was the standard, with a duration perceptually equivalent to an auditory duration of 500-ms. The auditory test duration varied across trials. The auditory test and visual standard were presented in mini-blocks of 10 trials with a fixed order of test and standard. Across mini-blocks the order of test vs. standard was randomized.

#### Main bimodal audiovisual conflict experiment

In the main task, both the standard and test stimuli included visual and auditory components. We introduced seven audiovisual duration conflicts, ranging from -250 to +250 ms (-250, -167, -83, 0, +83, +167, +250 ms) (Figure 3). We applied the conflict to both ends of the visual stimulus, i.e., half of the conflict magnitude extended the onset earlier and the other half extended the offset of the stimulus later. Likewise, half of the visual PSE bias, estimated based on the results of the cross-modal duration-comparison task, was subtracted from each end of the visual stimulus so that a nominally zero-conflict stimulus had a visual component that was perceived as having the same duration as the auditory component. Also, note that this bias correction was applied to both the standard and test stimuli, while the visual-duration perturbation was only added to the standard stimulus.

**Figure 3.**
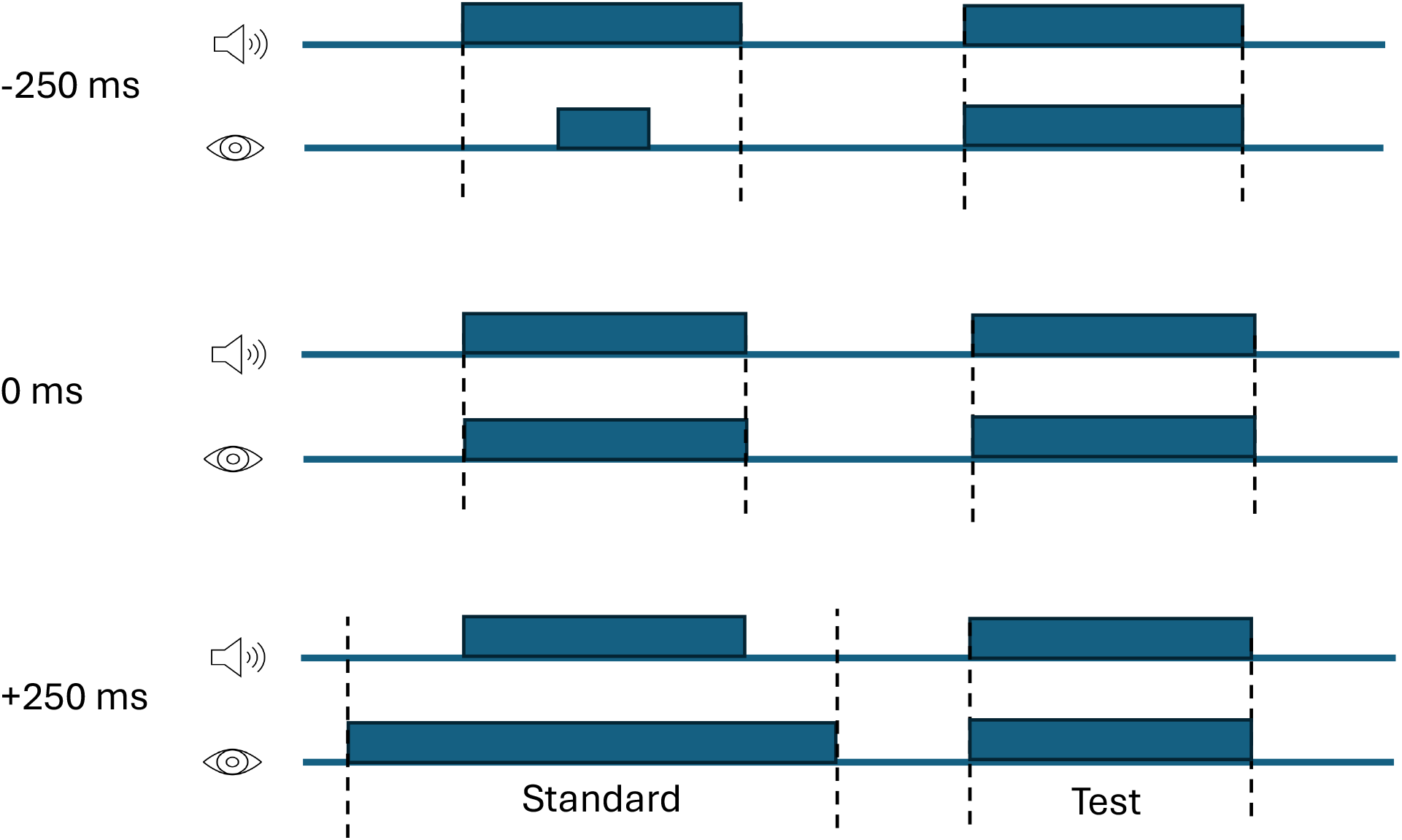
Audiovisual conflict in the bimodal experimental task. Conflicts were introduced exclusively in the visual component of the *standard* stimulus, while the auditory component remained unchanged. The top row shows an example negative-conflict (−250 ms), in which the visual duration of the standard stimulus was shorter than the auditory duration. Across trials, audiovisual conflict was randomly sampled from seven levels ranging from −250 ms to +250 ms. Note that the visual durations in both the standard and test stimuli were also corrected for the participant’s audiovisual bias as estimated from the cross-modal task.

Participants were asked to judge which interval had a longer duration based *only* on the auditory stimulus. Although participants were asked to judge the auditory stimulus, at the beginning of the experiment they were instructed to pay attention to both modalities and to maintain gaze at the screen even though they were not judging visual duration. We also included an audio icon to indicate that they needed to answer based on the auditory stimulus.

Each participant completed three sessions with different ranges of audiovisual conflict: (0, −167 and 250 ms), (−83 and 167 ms), and (−250 and 83 ms). The order of sessions was counterbalanced across participants. As in the unimodal and cross-modal experiments, easy lapse-rate trials were included in each condition (14 trials for each combination of auditory-noise level and cue conflict). For each condition (noise level and cue conflict), participants completed two interleaved staircases (one 1-down–2-up and one 2-down– 1-up), each consisting of 70 trials. Across the 7 conflict levels and 2 auditory-noise conditions, this resulted in a total of 2,156 trials for the main task.

#### Staircases

In all tasks, we measured discrimination thresholds using adaptive staircases, as described in each task section. Step-size reduction was applied after each staircase reversal. Specifically, whenever the staircase changed direction (from increasing to decreasing test duration, or vice versa), the step size was halved. Step-size reduction was applied only for the first two reversals; subsequent steps remained fixed.

The initial step size in the staircase procedure was defined as the duration difference percentage (Δ) between the test and standard durations. At the beginning of the experiment, Δ was set to the largest negative value allowed, corresponding to −95% of the standard duration. Since the standard duration was 0.5 s, this yielded an initial test interval of 0.5 − 0.95 *×* 0.5 = 0.025 s. This value represents the shortest possible test interval used in the experiment.

The starting value of the staircase (Δ) was set such that the resulting change in duration corresponded to approximately 475 ms for both auditory and visual cues. This ensured that the initial difference was clearly perceptible for participants.

### Data analysis

The log-normal observer model is well suited for duration judgments, as it naturally captures scalar variability consistent with Weber’s law and constrains duration estimates to be positive (Haigh et al., 2021). The psychometric function included three parameters: a lapse rate (*λ*), capturing stimulus-independent response errors, a sensory-noise parameter (*σ*_*v*_ for the visual task and *σ*_*a,l*_ and *σ*_*a,h*_ for the low and high auditory-noise conditions), reflecting the uncertainty in sensory encoding in log-duration space (a smaller value of *σ* corresponds to a steeper psychometric function) and a bias term (*µ*).

Under this model, the probability of judging the test interval as longer than the standard is given by

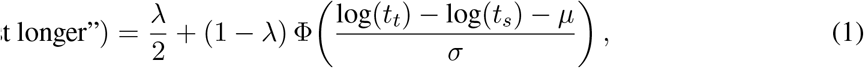

where *t*_*t*_ and *t*_*s*_ denote the test and standard durations, respectively, and Φ is the standard normal cumulative distribution function.

We did not include a bias parameter (i.e., *µ* = 0) for fitting the unimodal experiments. In a symmetric two-interval, forced-choice (2IFC) design, any additive bias in perceived log-duration would affect both the standard and test intervals equally and therefore cancel in the comparison. Consequently, when the standard (*t*_*s*_) and the test (*t*_*t*_) durations are identical, the model predicts *P* (“test longer”) = 0.5 by symmetry, and the bias term is inappropriate.

We conducted all analyses on the raw log-space sensory-noise estimates (*σ*_*v*_, *σ*_*a,l*_ and *σ*_*a,h*_). However, for reporting purposes we converted these to just noticeable differences (JNDs) in milliseconds, defined as the minimum duration increment required to reach 75% on the lapse-free component of the fitted psychometric function, and then to Weber fractions. Thus, although the lapse rate (*λ*) was estimated in the psychometric fits, the *σ*-to-JND conversion below reports lapse-corrected sensory precision. Given a fitted value of *σ*, the JND at this lapse-free 75% criterion is:

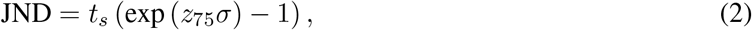

where *z*_75_ = Φ^−1^(0.75) is the inverse cumulative normal value corresponding to the 75% point of the lapse-free cumulative normal. We then computed the Weber fraction as the proportional threshold:

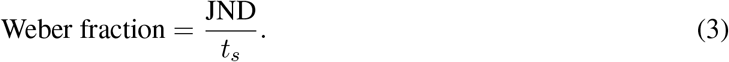

## Results

### Cue reliability

To measure sensory precision, all participants completed two unimodal duration-discrimination tasks: one auditory and one visual (Figure 1; see Methods for details). For each task, we fit log-normal psychometric functions to individual participants’ data using maximum-likelihood estimation. Figure 4 shows the psychometric-function fits (Panel A) for the auditory (high/low noise) and visual conditions and the unimodal duration-discrimination thresholds (Weber fraction; Panel B).

**Figure 4.**
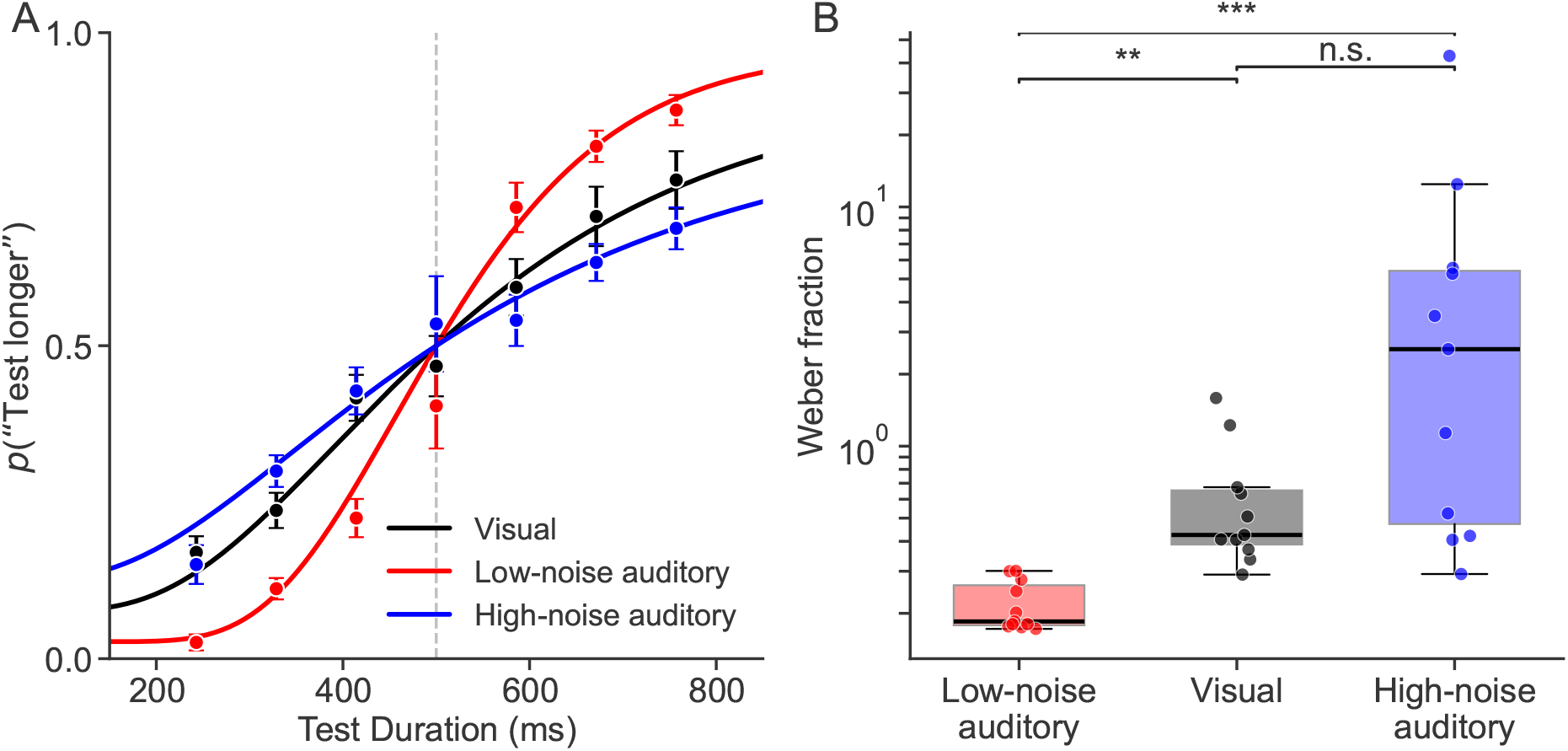
Unimodal visual and auditory duration-discrimination psychometric functions. (**A**) Psychometric functions showing *p*(“test longer”) as a function of test duration, pooled across all participants and trials. Data points represent proportions calculated within duration bins; error bars indicate binomial standard errors. Solid lines show fitted log-normal psychometric functions. Red: low-noise auditory; black: visual; blue: high-noise auditory. Discrimination precision was highest for low-noise auditory, intermediate for visual, and lowest for high-noise auditory conditions. (**B**) Unimodal discrimination thresholds expressed as Weber fractions, derived from log-normal psychometric fits, for the low-noise auditory (red), visual (black), and high-noise auditory (blue) conditions. Each point represents an individual participant. Boxplots display the interquartile range (IQR; 25th–75th percentile), with the median indicated by the central line. Whiskers extend to the most extreme values within 1.5 *×* IQR of the first and third quartiles; values beyond this range are treated as outliers. Data points are horizontally jittered for visualization clarity.

The Friedman test and Wilcoxon signed-rank test analysis revealed significant effects of the auditory noise level on duration-discrimination precision. For the low- and high-noise auditory and visual conditions, the estimated Weber fractions were 0.18, 2.54, 0.42, respectively, with corresponding median JNDs of 92.5, 1271.8, and 212.7 ms. A Friedman test revealed a significant difference across conditions (*χ*^2^(2) = 18.73, *p <* .001; Figure 4), indicating that discrimination thresholds differed across the auditory (low- and high-noise) and visual conditions.

Pairwise comparisons were conducted using Wilcoxon signed-rank tests with Bonferroni correction (*α* = .0167). *σ*_*a,l*_ (low auditory-noise condition) was significantly lower than *σ*_*a,h*_ (high auditory-noise condition; *W* = 0, *p <* .001). The value *W* = 0 indicates that, for every participant, the paired difference *σ*_*a,h*_ − *σ*_*a,l*_ was positive, i.e., *σ*_*a,l*_ was lower than *σ*_*a,h*_ for all participants. Moreover, *σ*_*a,l*_ was significantly lower than *σ*_*v*_ (*p <* .001). The difference between *σ*_*v*_ and *σ*_*a,h*_ was not significant after Bonferroni correction (*p* = 0.18).

### Modality-specific duration biases

To estimate participants’ inherent modality-specific biases in duration perception, we included a cross-modal duration-comparison task (Figure 2). In this experiment, participants were presented with an auditory signal (embedded in continuous auditory noise) and a visual signal (a filled disk against an otherwise empty background). The presentation order of these stimuli was precued at the beginning of each mini-block of 10 trials. Participants reported which stimulus had the longer duration.

Psychometric fits revealed systematic biases in cross-modal judgments, with bias magnitude depending on auditory noise level (Figure 5). Under the low auditory-noise condition, the median point of subjective equality (PSE) shift was −116 ms, indicating that the auditory duration had to be physically shorter than the visual duration to be perceived as equal. A Wilcoxon signed-rank test confirmed that this shift differed significantly from zero (*W* = 6, *p* = .014). Under the high auditory-noise condition, the median PSE shift was −73 ms, indicating a reduced auditory bias. However, this shift did not differ significantly from zero. Together, these results indicate a reliable overestimation of auditory relative to visual duration in the low-noise condition, with a weaker bias under high auditory noise.

**Figure 5.**
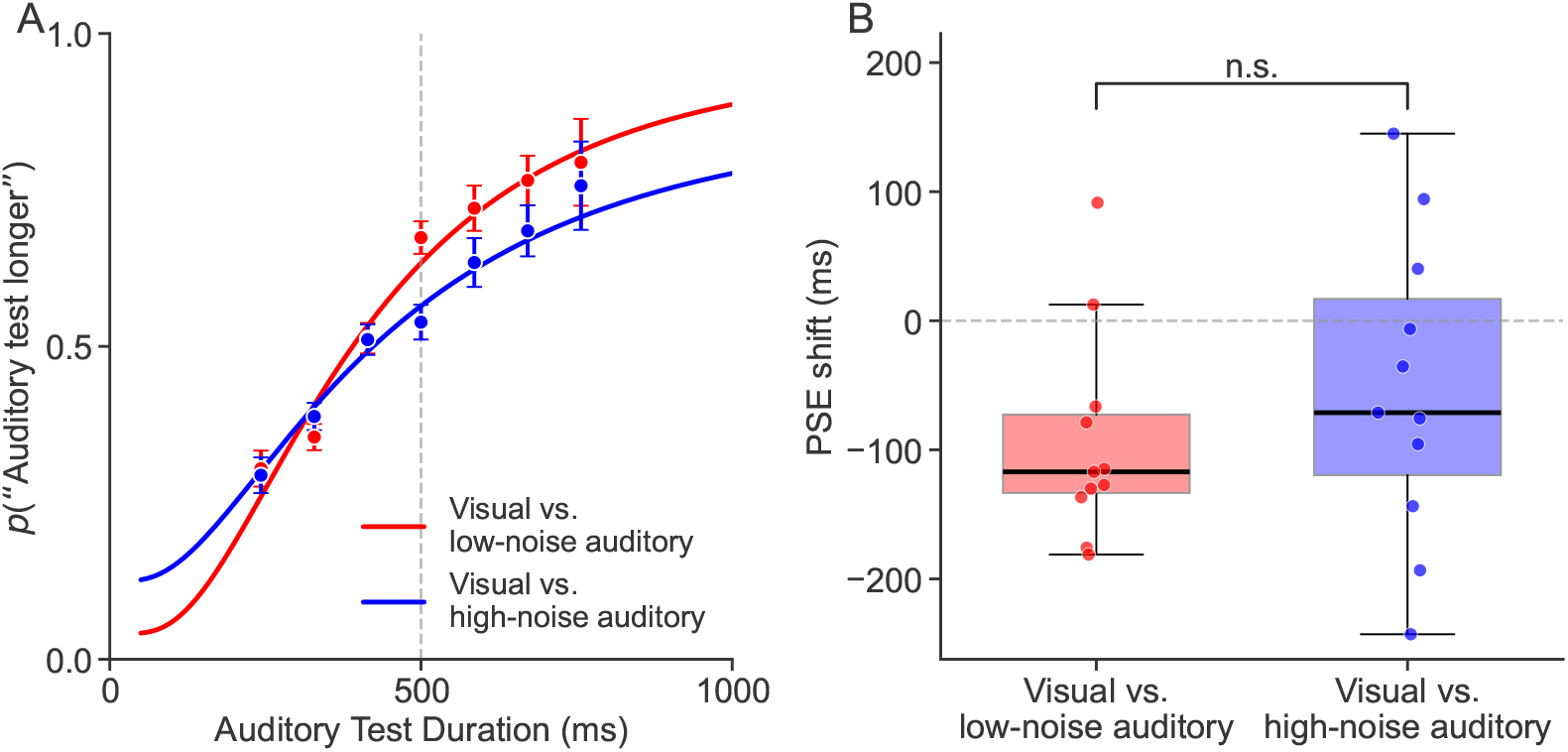
PSE shifts and psychometric curve fits for the cross-modal duration comparison. (**A**) Psychometric functions (dots) and model fits (solid lines), showing *p*(“auditory test longer”). Binary data were grouped into 7 equally-spaced bins between 0 and 1000 ms (error bars: SEM). For this plot we pooled the data across all participants then fit with log-normal cumulative distribution functions (parameters: *λ*: lapse rate; *µ*: PSE, *σ*: log-space noise SD). Solid lines show fitted log-normal psychometric functions. (**B**) Point of subjective equality (PSE) shifts for the low-noise (red) and high-noise (blue) auditory conditions. Superimposed are the individual participant data points (horizontally jittered). The PSE shift is the difference between the standard duration (500 ms) and the auditory duration that is perceived as identical to the standard.

Given the systematic modality-specific biases observed in the cross-modal task, we estimated each participant’s subjective auditory–visual duration bias and compensated for it in the bimodal experiment. Specifically, we adjusted the visual duration by adding half of the measured bias to the onset and half to the offset of the visual duration. This procedure preserved audiovisual synchrony while ensuring that, in the absence of added conflict, auditory and visual components were perceptually matched in duration.

### Main experiment: Cue-conflict effects

In the main bimodal experiment, participants were asked to judge which of two stimulus intervals was longer based solely on the auditory duration (Figure 3). Despite this instruction, participants’ judgments were systematically influenced by the duration of the visual stimulus, especially under high auditory noise.

To investigate how participants combined auditory and visual cues during duration judgments, we analyzed psychometric functions across varying levels of audiovisual conflict and auditory cue reliability. Figure 6 shows the psychometric functions for each conflict level, separately for the low (left) and high (right) auditory-noise conditions. As the visual duration increased, the curves shifted horizontally toward longer durations, indicating that the task-irrelevant visual cue biased auditory duration judgments. This bias was more pronounced under the high-noise condition, qualitatively consistent with the predictions from reliability-weighted cue combination. Under low noise, psychometric functions showed smaller shifts, reflecting a stronger reliance on the auditory modality.

**Figure 6.**
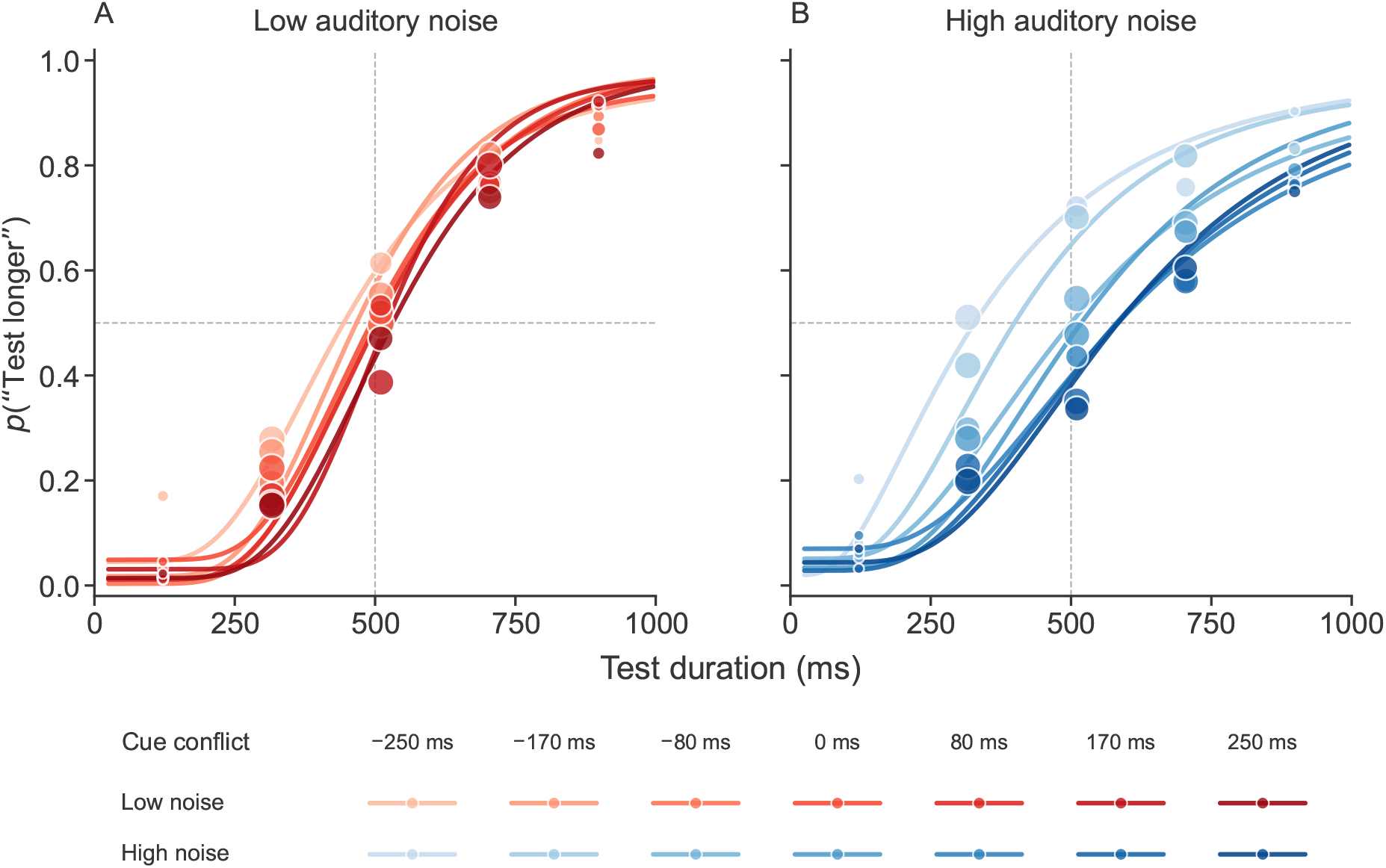
Pooled trial data across participants and fitted psychometric functions in the bimodal experiment. (A) Fitted psychometric functions in the low auditory-noise condition. Each curve corresponds to one of seven audiovisual conflict levels (−250 to +250 ms). Dots show *p*(“test longer”) as a function of seven equally spaced test-duration bins. Data points are plotted at the centers of the bins, with dot diameter proportional to the number of trials within each bin. (**B**) Same as (A), but for the high auditory-noise condition. Psychometric curves shift progressively with conflict magnitude, indicating stronger visual influence on perceived auditory duration under high auditory noise.

To quantify the results, we extracted the PSE from each psychometric curve and plotted it as a function of the conflict level (Figure 7). Under the high auditory-noise condition, PSEs showed a nearly linear relationship with conflict magnitude. In contrast, under the low-noise condition, PSEs stayed close to zero, indicating reduced bias toward visual durations. The error bars reflect the standard error of the mean across the 11 participants. We quantified this effect by fitting a linear regression of PSE shift on signed cue-conflict level separately for each participant and auditory-noise condition. The resulting slopes were positive in both conditions (low noise: mean slope = 0.148 ms/ms, SEM = 0.048, *t*(10) = 3.11, *p* = .011; high noise: mean slope = 0.505 ms/ms, SEM = 0.083, *t*(10) = 6.11, *p <* .001). Slopes were significantly larger under high than low auditory noise (paired *t*(10) = 5.65, *p <* .001), indicating that auditory duration perception was systematically biased toward the visual duration, and that the strength of this bias was modulated by auditory cue reliability.

**Figure 7.**
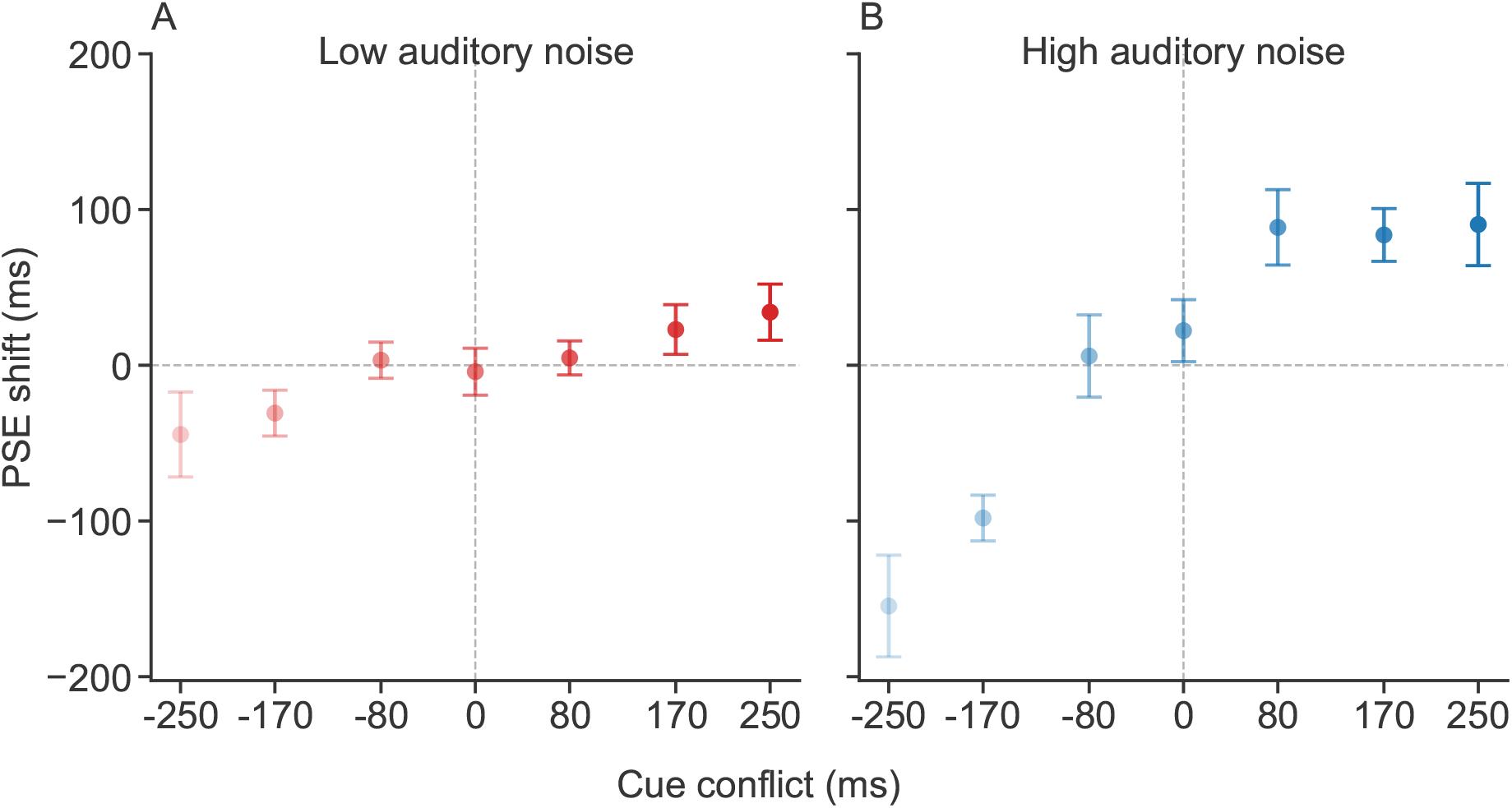
Effect of audiovisual conflict on perceived auditory duration. Group-averaged values of the shift of the point of subjective equality (PSE) toward the visual duration as a function of audiovisual duration conflict, shown separately for the low-noise (**A**) and high-noise (**B**) auditory conditions. Each data point reflects the mean PSE shift across 11 participants; error bars show SEM across participants. Under low noise, PSEs are more stable across conflict levels, suggesting auditory dominance. Under high noise, auditory duration judgments are pulled toward the visual duration.

## Computational models

To explain participants’ behavior in the audiovisual duration-conflict task, we fit three models to the behavioral data. These models share the same encoding process but differ in decoding strategy. Below, we describe the models’ encoding process, followed by the decoding stage.

### Sensory encoding

All models assume that internal representations of auditory and visual durations are noisy. To capture the Weber-like scaling of temporal uncertainty (scalar variability), we assume that durations are encoded in logarithmic space with additive Gaussian noise:

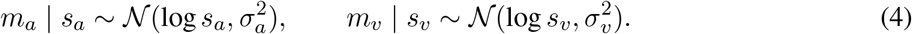

Here, *s*_*a*_ and *s*_*v*_ denote stimulus durations in linear time units, whereas *m*_*a*_ and *m*_*v*_ represent internal measurements in log-duration space. Because encoding noise is Gaussian in log space, the implied distribution of internal measurements expressed in linear time is log-normal. This formulation naturally produces scalar variability, such that uncertainty scales proportionally with stimulus duration. Scalar variability is a characteristic feature of perception of time and motivates this log-duration encoding. Thus, both encoding and decoding are performed in log-duration space (Killeen and Weiss, 1987; Wittmann, 2009; Ren et al., 2020).

### Decoding strategy

While all tested models share the same encoding process, they differ in how encoded information is integrated and read out across sensory modalities. We consider three classes of integration and decision strategies: forced fusion, causal inference, and heuristic cue selection. We describe each of the three models next.

### Forced fusion (Optimal cue integration)

The forced-fusion model assumes that observers integrate auditory and visual signals for the auditory-duration judgment regardless of cue-conflict magnitude. Moreover, the integration is optimal, with each cue weighted by its reliability (Eq. 23).

This model can be viewed as assuming that the probability that the two cues originate from the same source is 100%, so cues are always integrated regardless of cue-conflict magnitude.

### Causal-inference model

The Bayesian causal-inference model assumes that the observer first infers whether the auditory and visual cues arise from a common source by computing the posterior probability of a common source given the two cues’ sensory measurements:

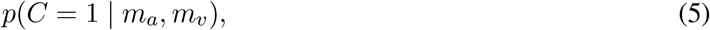

where *C* = 1 denotes a common cause. The observer then forms a posterior over the task-relevant variable (here, the auditory stimulus duration *s*_*a*_) by averaging over two posteriors. The observer’s posterior distribution over auditory duration is given by:

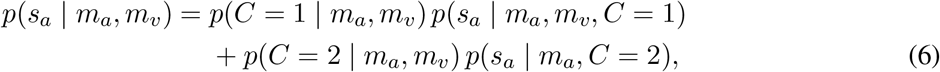

in which *p*(*C* = 2|*m*_*v*_, *m*_*a*_) indicates the posterior probability of separate causes. *p*(*s*_*a*_ | *m*_*v*_, *m*_*a*_, *C* = 1) and *p*(*s*_*a*_ | *m*_*a*_, *C* = 2) specify the posteriors over *s*_*a*_ under each causal scenario. The posterior probability of common cause is computed as:

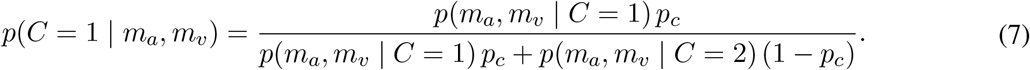

This model includes the prior probability of common cause, *p*_*c*_, which is fitted to the data. Likelihoods under the common-cause and separate-causes hypotheses are computed by marginalizing over the latent true stimulus durations using a bounded prior, following the standard formulation of Bayesian causal inference (Körding et al., 2007). Full expressions, choice of prior distribution, and analytical derivations are provided in the Appendix, specifically in the Bayesian causal-inference model subsection.

Let *s*_min_ and *s*_max_ denote the lower and upper physical-duration bounds used by the model. Because inference is performed in log-duration space, the model uses a bounded uniform prior over [*t*_min_, *t*_max_], where *t*_min_ = log *s*_min_ and *t*_max_ = log *s*_max_. These bounds provide finite support for the latent log-duration variable.

Here, the model assumes that participants infer the stimulus duration based on the inferred causal relationship between the cues presented in the trial. The observer first infers the posterior probability that both cues originate from a common cause by computing *p*(*C* = 1 | *m*_*a*_, *m*_*v*_). Under this model, observers compute two estimates of the auditory stimulus duration, one based on the common-cause scenario that relies on both the auditory and visual measurements, and another for the separate-causes scenario based on the auditory measurement alone. For each scenario, the model uses the Bayes least-squares (BLS) estimate, that is, the posterior mean under squared-error loss. The final auditory duration estimate is a weighted average of these two estimates, given by:

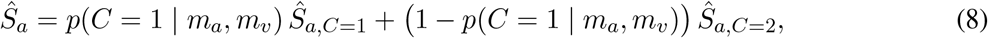

where *Ŝ*_*a,C*=1_ and *Ŝ*_*a,C*=2_ denote the posterior-mean estimates of the auditory stimulus under the common-cause and separate-causes assumptions, respectively. Because the latent-duration prior has bounded uniform (boxcar) support, the posterior distributions are truncated Gaussian distributions. To clearly show the effects of the prior, we assumed a Bayes least-squares observer (using the mean of the posterior as the estimate) rather than a maximum a posteriori estimator (see Appendix for details). The above computation yields an internal estimate of auditory duration for a single interval. In the present two-interval forced-choice task, the same inference process is applied independently to the standard and test intervals, producing estimates *Ŝ*_std_ and *Ŝ*_test_. The observer then compares the estimated durations of the test and standard intervals and responds “test longer” whenever

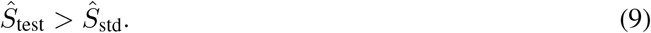

From the modeler’s perspective, the predicted response probability is obtained by marginalizing over the noisy sensory measurements:

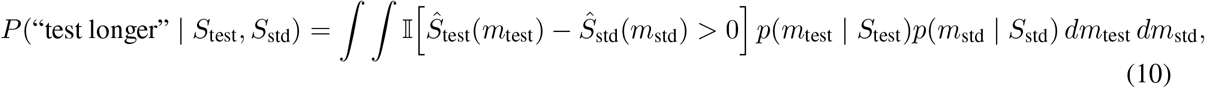

where I[·] denotes the indicator function.

In practice, this probability is approximated using Monte Carlo simulation by repeatedly sampling noisy measurements for the test and standard intervals, applying the model-specific decoding rule, and computing the proportion of samples for which the decoded test duration exceeded the decoded standard duration.

### Probabilistic cue switching

We also considered a probabilistic cue-switching model, in which observers based their decision on either the auditory or the visual cue on each trial (see Methods). Instead of computing posterior probabilities of causal structure, this heuristic strategy selects a single cue stochastically. In auditory-noise condition *j* ∈ {*l, h*}, the decision is based solely on the visual estimate with probability *p*_*v,j*_ and solely on the auditory estimate with probability 1 − *p*_*v,j*_. Thus, the model includes separate visual-selection probabilities of selecting the visual cue, *p*_*v,l*_ and *p*_*v,h*_, for the low- and high-auditory-noise conditions, respectively. Despite its simplicity, the switching strategy can reproduce cue-integration–like patterns when responses are averaged across trials.

### Modeling results

We first evaluate how well the models accounted for the shifts in PSE as a function of audiovisual cue conflict across the two auditory noise levels. We compared observed PSE shifts with predictions from three models: forced fusion, causal inference, and probabilistic cue switching. Sensory-noise parameters were fitted using the data from the bimodal experiment.

Figure 8 shows model fits to the PSE-shift data. All three models capture the overall pattern that perceived auditory duration was systematically biased toward visual duration and that the bias increased approximately linearly with audiovisual conflict magnitude. All three models also predict an overall larger bias under higher auditory noise. This convergence suggests that our data may be limited in the ability to adjudicate between these models.

**Figure 8.**
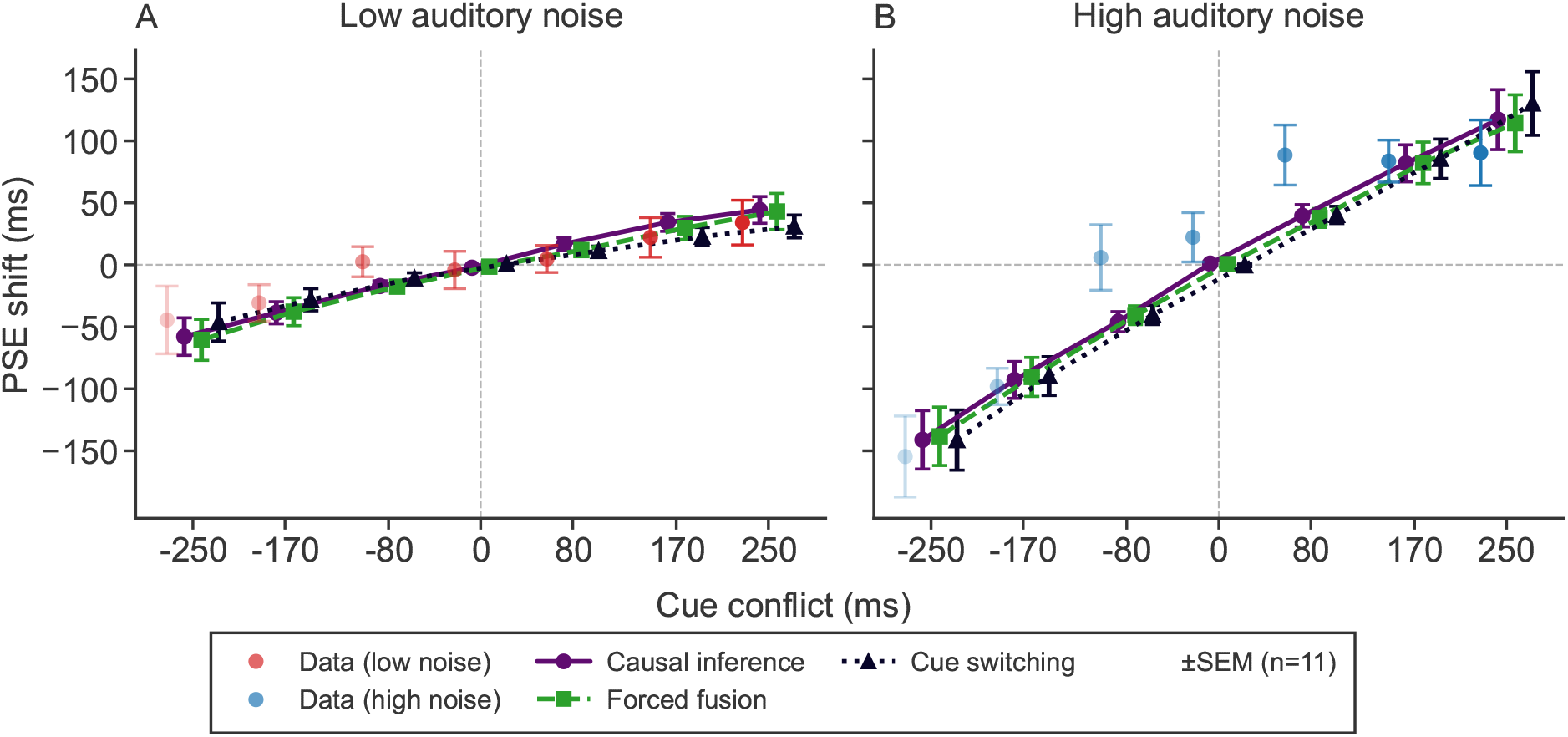
Measured and predicted shifts of the point of subjective equality (PSE) as a function of audiovisual conflict for the low auditory-noise condition (**A**) and high auditory-noise condition (**B**), averaged over 11 participants. Positive values indicate biases toward longer perceived auditory duration. Red and blue show empirical data. Other lines indicate model predictions for forced fusion, causal inference, and probabilistic cue switching. The model predictions are jittered horizontally for readability. All model predictions were generated using sensory-noise parameters fitted using the bimodal data. Error bars reflect *±* SEM across participants.

Causal-inference models typically predict a non-linear breakdown of integration for large cue conflicts. Our data show only weak deviations from linearity. These deviations are more pronounced for large positive conflicts than for large negative conflicts, indicating an asymmetry in bias magnitude across the conflict range. This pattern suggests that, at the group level, observers do not exhibit the strong nonlinearity predicted by the causal-inference model in which cue integration gradually changes to cue segregation with increases in cue conflict. Instead, integration remains approximately linear within the tested conflict range. As a result, the behavioral signatures of causal inference are weak, limiting our ability to discriminate between the causal-inference model and simpler integration or heuristic strategies with these data.

We also examined whether the models captured the sensitivity. The fit value of *σ* corresponds to the slope of the fitted psychometric functions, with larger values of *σ* (in log-duration units) corresponding to lower encoding precision and shallower slope. The probabilistic cue-switching model appeared to track the conflict-dependent changes in slope more closely than the forced-fusion and causal-inference models (Figure 9). To quantify this, we computed the root mean squared error (RMSE) between the measured *σ* values and those predicted by each model in both noise conditions. RMSE indeed tended to be lower for the cue-switching model (0.048 log units) than for the causal-inference model (0.066) or the forced-fusion model (0.072).

**Figure 9.**
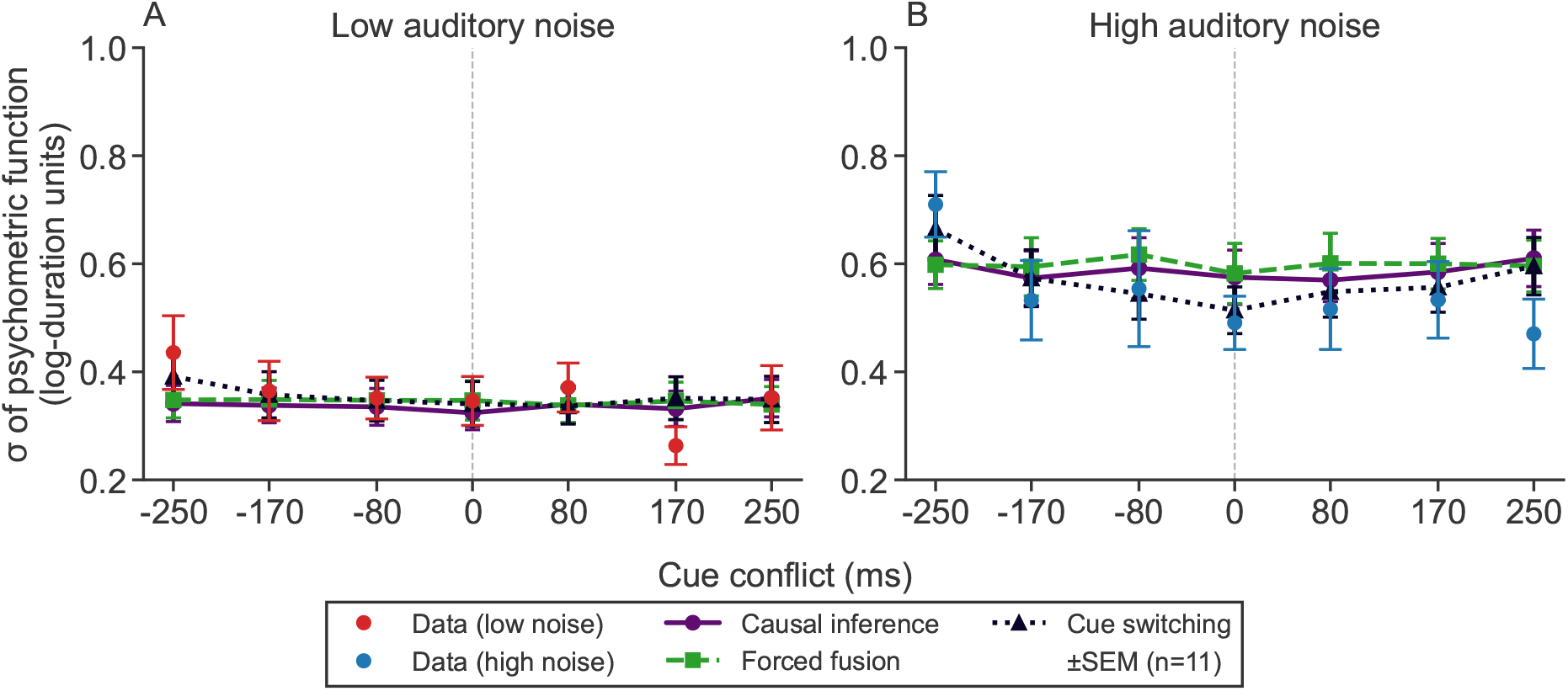
Measured and predicted overall noise (*σ*) parameters of psychometric functions. *σ* values are shown in log-duration units as a function of audiovisual conflict for the low (**A**) and high (**B**) auditory-noise conditions, averaged over 11 participants. Red and blue points show empirical data (group mean *±* SEM across participants). Other lines indicate model predictions for forced fusion, causal inference, and probabilistic cue switching. Error bars indicate SEM across participants. Larger *σ* values indicate lower duration-discrimination precision.

### Model fitting and model comparison

All models have a lapse-rate (*λ*) parameter, two auditory duration-noise parameters (one for each auditory-noise condition: *σ*_*a,l*_ and *σ*_*a,h*_), and one visual duration-noise parameter (*σ*_*v*_). For models that incorporate causal inference, there is an additional parameter (*p*_*c*_), which captures the prior probability of a common source. For the probabilistic cue-switching model, there are two additional parameters (*p*_*v,l*_ and *p*_*v,h*_) that specify the probability of switching (i.e., relying on vision rather than audition) in each of the two auditory-noise conditions.

We fit each model to each participant’s data (from the two auditory-noise conditions jointly) using maximum-likelihood estimation. For each model, we computed the predicted probability of responding “test longer” for every stimulus condition via Monte Carlo simulation. Specifically, on each simulated trial, we drew noisy internal measurements of auditory and visual durations from the encoding model, applied the model’s integration or decision rule to produce an estimate of the auditory stimulus, and then determined whether the test estimate was longer than the standard estimate. For each condition, that is, for each combination of auditory test duration, standard duration, cue conflict, and noise level, we simulated 3000 Monte Carlo trials (i.e., repeated draws of internal sensory measurements). For each Monte-Carlo trial we generated samples of the auditory and visual internal measurements according to the model’s encoding-noise assumptions and determined the model’s response for that trial. The predicted response probability for each condition was computed as the proportion of simulated trials for that condition for which the model observer judged the test duration as longer. The log-likelihood of each model was then computed using the binomial likelihood of the observed responses given the predicted probabilities across all conditions. Parameter bounds were chosen based on plausible ranges for sensory noise and lapse rates. The physical bounds *s*_min_ and *s*_max_ were fixed to the minimum and maximum test durations for each participant. Thus, the log-normal models applied the bounded uniform prior over the corresponding log-space interval [log *s*_min_, log *s*_max_]. Parameters were estimated by minimizing the negative log-likelihood using Bayesian Adaptive Direct Search (Acerbi and Ma, 2017). To reduce the risk of local minima, the optimization was repeated from 10 random initial parameter values.

Sensory-noise parameters were fitted using the data from the bimodal task, because effective noise may differ between unimodal and bimodal contexts due to attentional allocation and cross-modal interactions (Badde et al., 2020). To ensure robustness, we also evaluated all models using sensory-noise parameters estimated using the psychometric function fits to the unimodal-task data. The results using these parameter estimates were qualitatively consistent (Figure 15).

We evaluated model performance using the Akaike Information Criterion (AIC) at the individual participant level. Consistent with the observation that all models captured the data reasonably well (Figure 8), model comparison revealed only small differences in fit quality across integration strategies.

#### Fixed sensory noise

In this analysis, we assume that sensory noise is identical for unimodal and bimodal stimuli. Thus, we estimated the sensory-noise parameters using the data from the unimodal duration-discrimination tasks, and then used those estimates when fitting the data from the bimodal task. Figure 10A shows the participant-level ΔAIC values, computed as the difference between each model’s AIC for each participant and the AIC values of the Forced fusion model. For most participants (8 out of 11), the probabilistic cue-switching model achieved the lowest ΔAIC among the candidate models, indicating that it provided the best trade-off between goodness of fit and model complexity. The Bayesian causal-inference model was the best-fitting model for the remaining three participants. The forced-fusion model did not achieve the lowest ΔAIC for any participant. At the group level, the summed ΔAIC values across participants yielded the same pattern, with the probabilistic cue-switching model overall performing the best (Figure 10A right panel).

**Figure 10.**
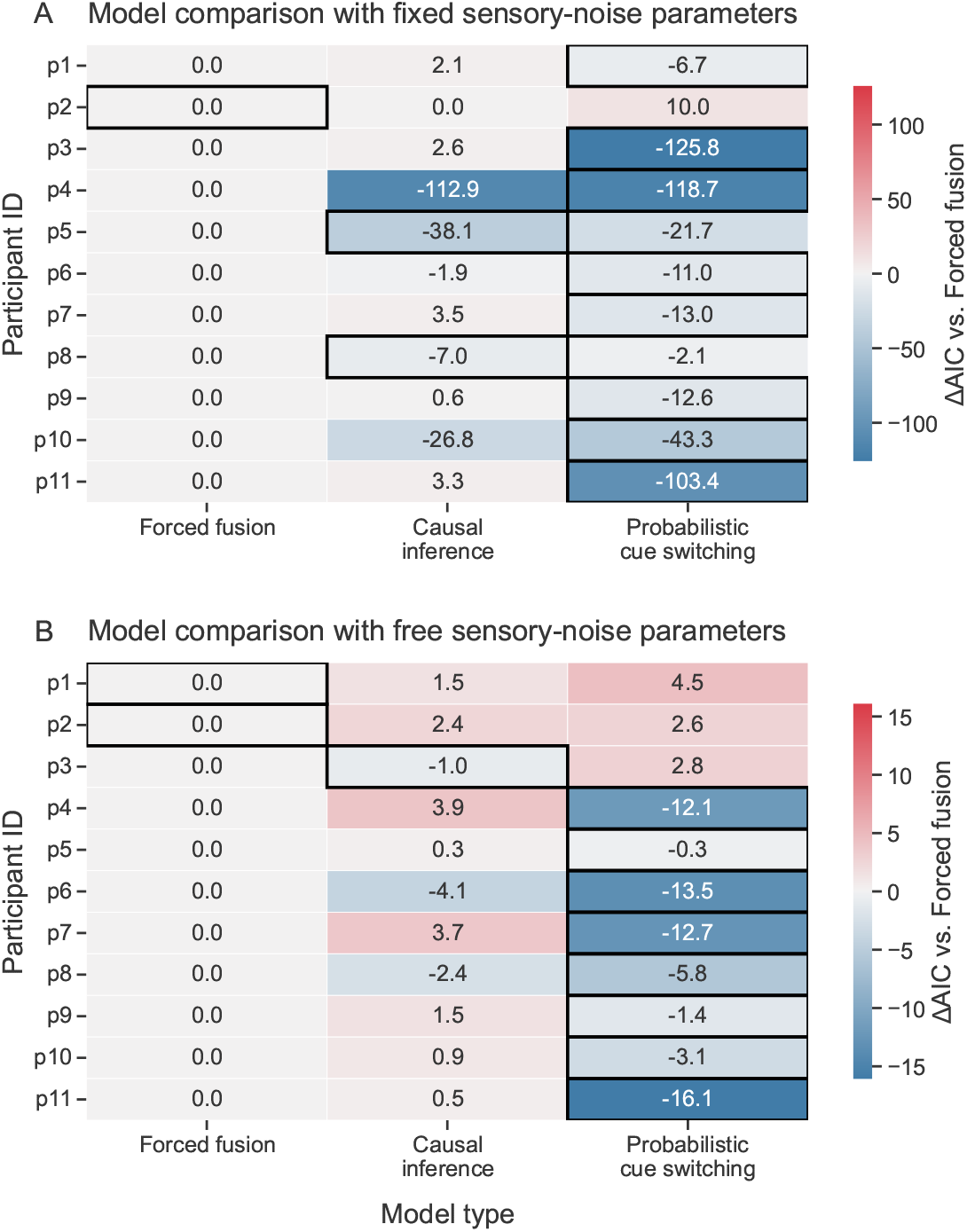
(**A**) Model comparison assuming sensory-noise parameters fixed to values estimated from the unimodal tasks. (**B**) Model comparison when sensory-noise parameters were fit to the bimodal data. Each panel shows participant-level ΔAIC values for each model, computed relative to the forced-fusion model (the baseline column, which is therefore identically zero); negative values (blue) indicate a better fit than forced fusion. Rows correspond to participants and columns correspond to models, and the best-fitting model (lowest AIC) for each participant is outlined. When sensory-noise parameters were fixed (A), the probabilistic cue-switching model provided the best fit for 8 of 11 participants. When the sensory-noise parameters were fit directly to the bimodal data (B), the cue-switching model provided the best fit for 8 of 11 participants.

#### Free sensory noise

When sensory-noise parameters were fitted using the bimodal data, model comparison revealed substantially weaker separation between candidate models (Figure 10B). Participant-level ΔAIC values showed considerable overlap between models, indicating that these integration strategies provided similarly good fits once the sensory noise was allowed to vary.

The probabilistic cue-switching model provided the best fits for the largest number of participants (8 out of 11; Figure 10B). However, the individual ΔAIC values were very low, suggesting limited discriminability between decoding strategies under this assumption.

#### Group-level model comparison and its reliability

To summarize model performance across participants, we summed each participant’s ΔAIC relative to the forced-fusion model: a negative sum means the model had a lower total AIC than forced fusion across the group, and the further below zero, the stronger the group-level preference (Figure 11).

**Figure 11.**
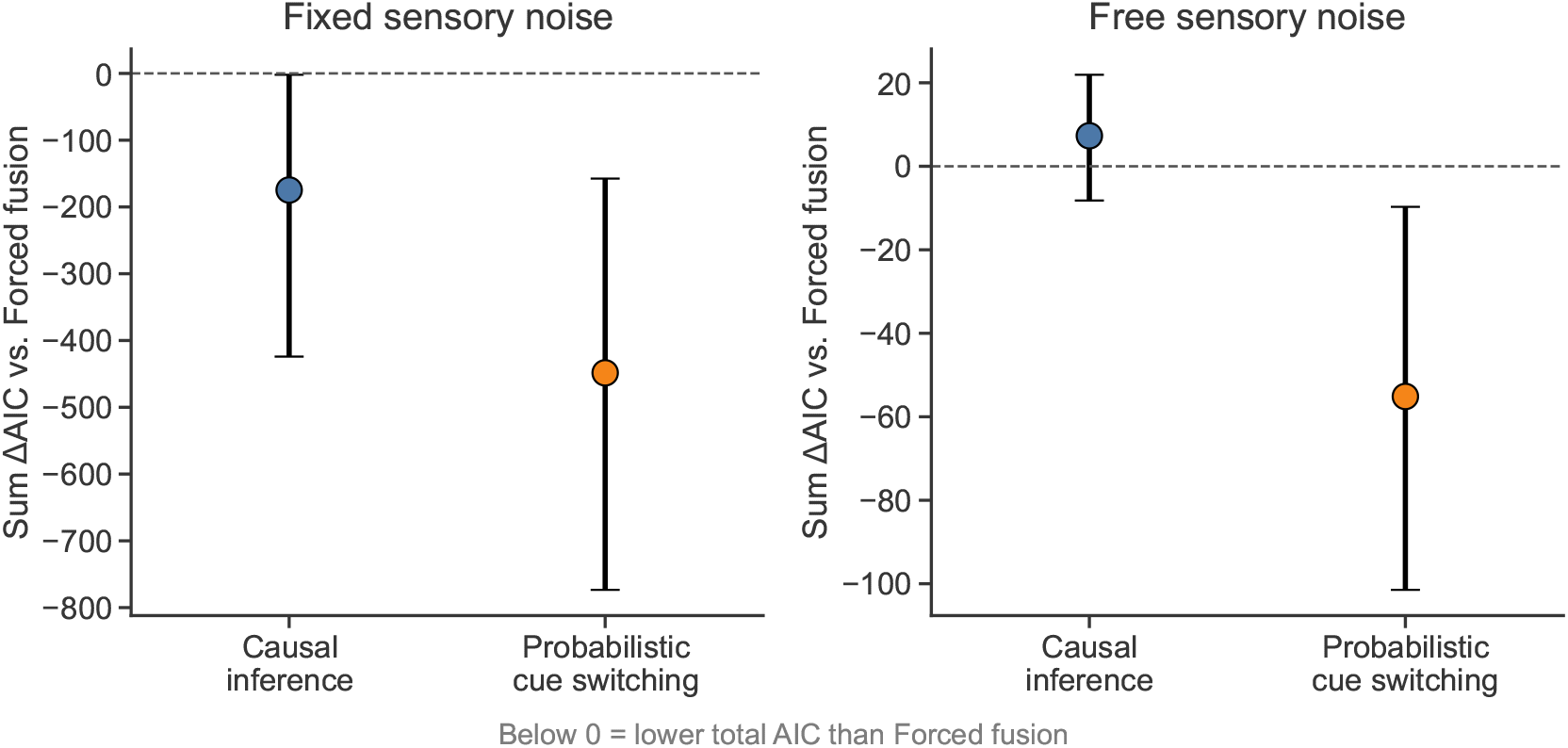
Group-level model comparison: summed ΔAIC. Sum of the per-participant ΔAIC relative to the forced-fusion model (the baseline, fixed at 0) under fixed (left) and free (right) sensory noise. Each colored point is the total across the 11 participants for one model, with 95% confidence intervals from 10,000 participant bootstrap resamples. Values below the dashed zero line indicate a lower total AIC, i.e. a better fit, than forced fusion. Under fixed sensory noise, the probabilistic cue-switching and causal-inference models fit clearly better than forced fusion (intervals below zero); under free sensory noise the totals are much smaller and the confidence intervals overlap zero.

Under fixed sensory noise, both the probabilistic cue-switching model (summed ΔAIC = −448, 95% CI [−764, −173]) and the causal-inference model (−204, [−471, 28]) fit better than forced fusion, with cue switching preferred overall; both confidence intervals fell entirely below zero. Under free sensory noise, the same ordering held, but the totals were much smaller and no longer reliable (cue switching −34, [−73, +1]; causal inference −1, [−17, +14]), with both confidence intervals overlapping zero.

This lack of separation under free sensory noise is not an artifact of Monte-Carlo evaluation noise. The per-participant AIC noise floor (median SD 1.9 AIC points) was comparable in magnitude to the per-participant ΔAIC itself, so individual-participant model assignments are unreliable in this regime; averaging across the 11 participants, however, reduced the Monte-Carlo contribution to below 1 AIC point. The null result therefore reflects genuine between-participant variability in model preference rather than simulation noise.

We complemented these criteria with random-effects Bayesian model selection, which estimates the frequency of each model in the population and is robust to outlying participants (Stephan et al., 2009; Rigoux et al., 2014). Probabilistic cue switching was the most frequent model under both assumptions about sensory noise, with an estimated model frequency of 0.65 and an exceedance probability of 0.93 under fixed sensory noise, and 0.46 (exceedance probability 0.68) under free sensory noise. Consistent with the small ΔAIC differences, however, the Bayesian omnibus risk was high (0.87 and 0.77, respectively), indicating that the candidate models cannot be confidently separated at the group level; the protected exceedance probabilities were correspondingly modest (approximately 0.41 for cue switching in both analyses).

#### Switching probabilities increase with auditory noise

Beyond model comparison, the parameters of the favored model were themselves consistent with the cue-switching account. The fitted probability of relying on the visual duration (*p*_*v*_) increased markedly with auditory noise, rising from a group mean of 0.25 under low auditory noise to 0.62 under high auditory noise (Wilcoxon signed-rank test, *p* = 0.001; an increase in 11 of 11 participants; Figure 12). Thus, observers were estimated to rely on the visual duration on roughly one-quarter of trials when auditory noise was low and nearly two-thirds of trials when it was high, providing a concrete and consistent mechanism for the noise-dependent visual bias.

**Figure 12.**
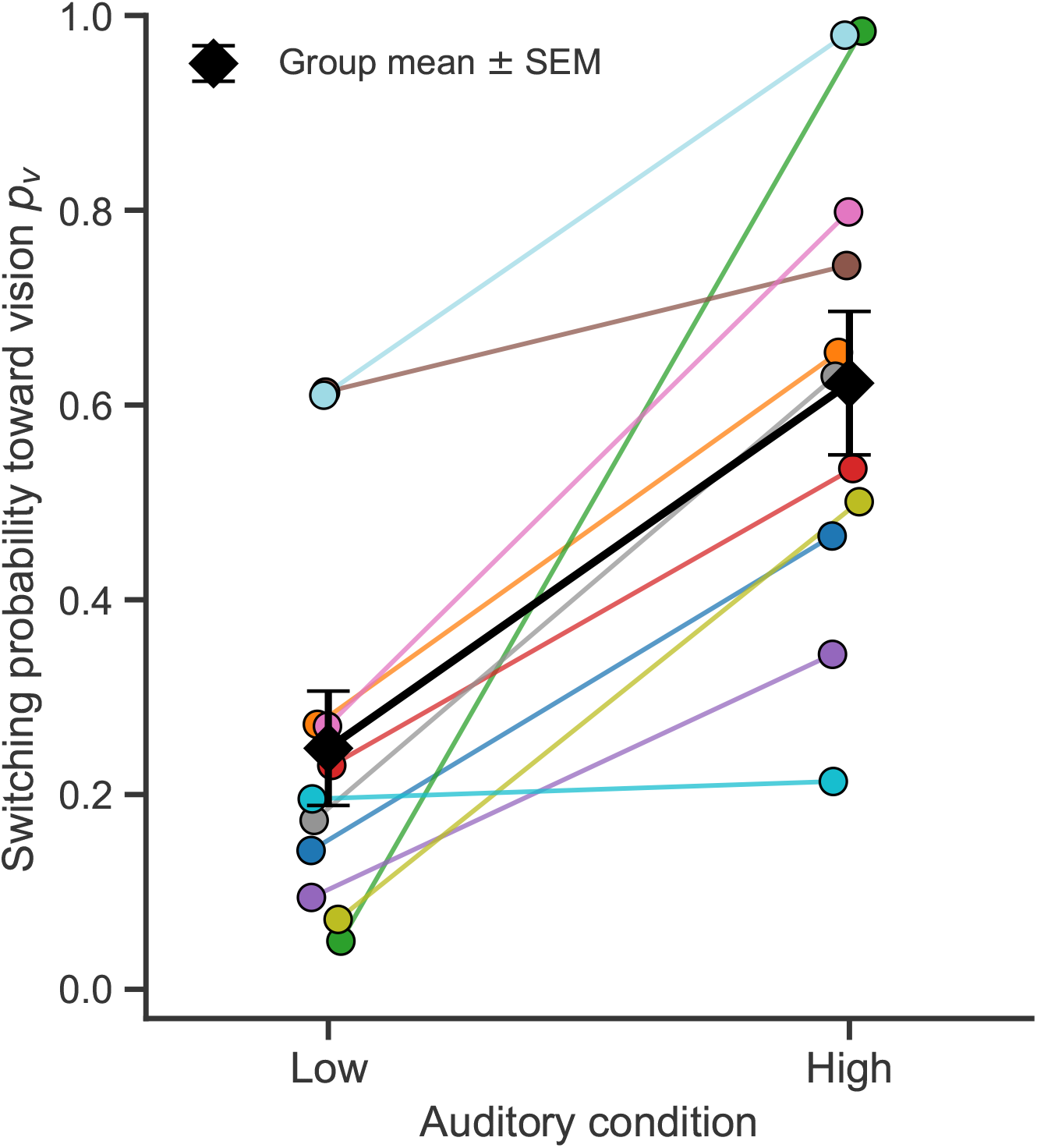
Fitted switching probabilities increase with auditory noise. Each colored thin line shows one participant’s fitted probability of relying on the visual duration (*p*_*v*_) in the low- and high-auditory-noise conditions, estimated from the probabilistic cue-switching model (free sensory noise fits). The probability of relying on vision increased with auditory noise in all participants (group means 0.25 → 0.62; Wilcoxon signed-rank *p* = 0.001). Black diamonds and line show group means *±* SEM.

#### High sensory-encoding noise limits model discriminability

To explain the lack of evidence for causal inference even at large audiovisual cue conflicts, we evaluated the posterior probability of a common cause *P* (*C* = 1 | *m*_*a*_, *m*_*v*_) across conflict levels using each participant’s fit parameters.

Figure 13 shows *P* (*C* = 1 | *m*_*a*_, *m*_*v*_) for an example participant. Across all conflict levels and auditory-noise conditions, the posterior probability of common cause remained relatively high (i.e., above 0.5), a pattern that was maintained for all participants (Figure 16). This is due to the large sensory encoding noise estimated from the data. Specifically, when sensory encoding noise is high, even substantial physical conflicts produce overlapping measurement distributions, yielding consistently high posterior probabilities of a common cause. This indicates that the observer’s fitted parameters are consistent with treating the auditory and visual signals as originating from a common source regardless of the imposed conflict.

**Figure 13.**
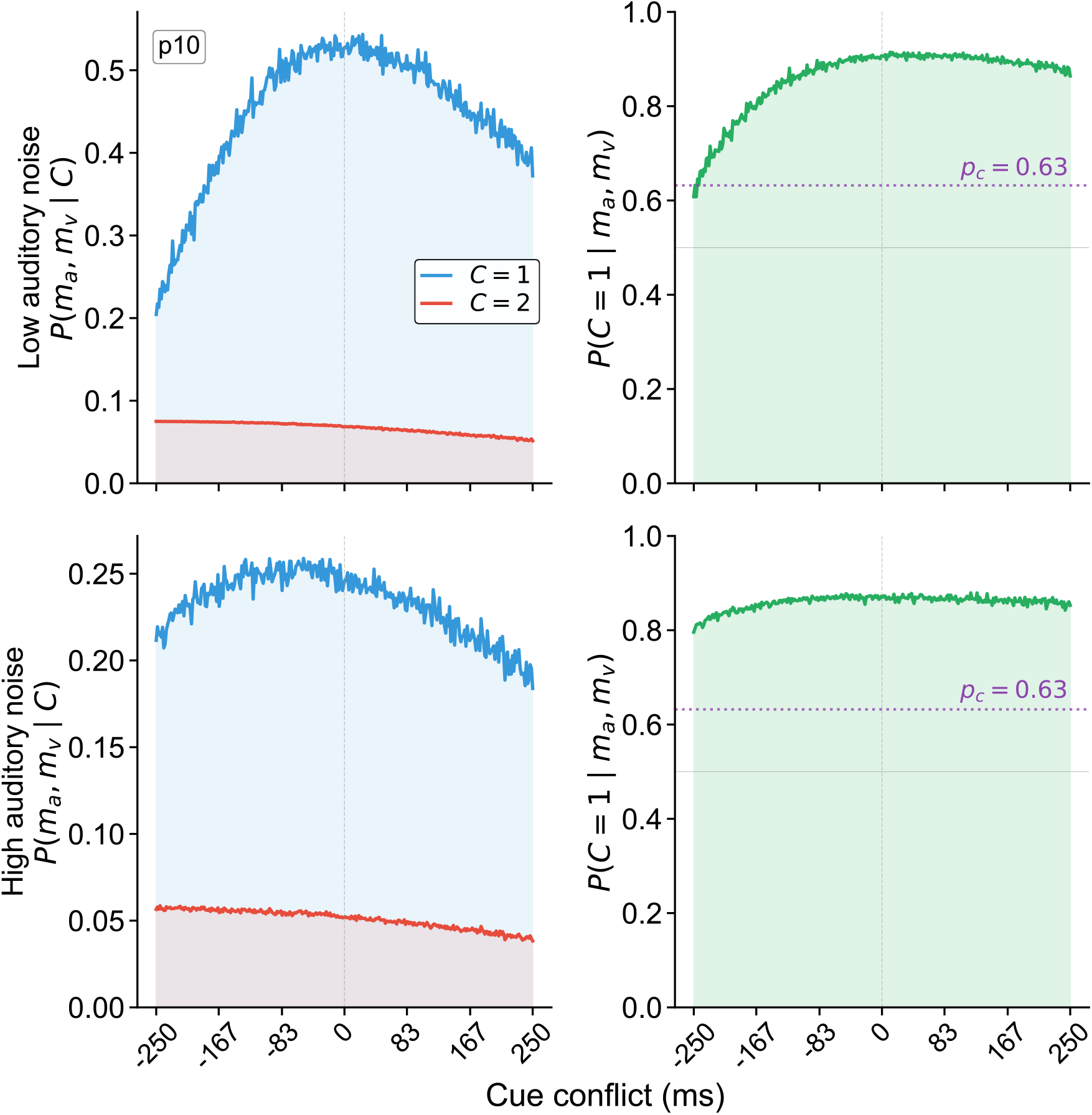
Posterior probability of a common cause across conflict levels. Likelihoods and posterior probability of a common cause for an example participant, plotted as a function of audiovisual duration conflict. Top row: low-auditory-noise condition; bottom row: high-auditory-noise condition. Left column: likelihood of the measurements under a common cause, *P* (*m*_*a*_, *m*_*v*_ | *C*=1) (blue) and under separate causes, *P* (*m*_*a*_, *m*_*v*_ | *C*=2) (red), plotted on the same axes so that their relative magnitudes can be compared directly. Right column: Posterior probability of a common cause, *P* (*C*=1 | *m*_*a*_, *m*_*v*_), with the dotted line marking the fitted prior *p*_*c*_. All quantities were computed via Monte Carlo simulation (500 samples per conflict level) using participant-specific sensory-noise estimates (*σ*_*a,l*_, *σ*_*a,h*_, *σ*_*v*_) and common-cause prior (*p*_*c*_) obtained from the log-normal causal-inference model fit.

This analysis reveals why the causal-inference and forced-fusion models produced similar predictions in our experiment. Given the estimated sensory-noise parameters, the posterior probability of a common cause never dropped low enough to produce substantial segregation of cues. The observer’s inference remained dominated by the common-cause hypothesis across the entire conflict range, resulting in near-linear, fusion-like behavior. Within the conflict range examined here, competing models predict largely overlapping, noisy psychometric patterns, imposing a fundamental limit on model identifiability that is not due to optimization failure or Monte Carlo noise.

## Discussion

In this study, we investigated how humans infer duration under conflicting auditory and visual cues with varying levels of sensory uncertainty. Previous work has shown that audiovisual duration cues can be integrated in a statistically optimal, reliability-weighted manner (Shams et al., 2005; Hartcher-O’Brien et al., 2014). Extending this framework, we asked whether such integration is flexibly modulated by causal inference, as observed in audiovisual spatial-localization tasks where large cue conflicts induce a non-linear breakdown of integration (Körding et al., 2007; Badde et al., 2021; Hong et al., 2021). Contrary to this expectation, we found little evidence for a pronounced non-linear reduction in integration strength. Instead, auditory duration judgments remained largely linear and noise-dependent even at relatively large conflicts, suggesting that short sub-second duration judgments may rely more strongly on heuristic integration strategies than on inference about causal structure.

To interpret these findings, we compared a set of models spanning Bayesian causal inference and probabilistic cue-switching strategies. Although a probabilistic cue-switching model provided the best overall fit under several model-comparison criteria, forced-fusion and causal-inference models produced highly similar predictions within the experimentally plausible conflict range. While these candidate models differ substantially in their computational assumptions, their behavioral predictions converge under the levels of sensory conflict and uncertainty tested here. This convergence highlights a fundamental limitation of model identifiability. Within the conflict range tested here (*±*250 ms), and in the absence of externally added visual noise (i.e., visual stimuli with clear onset and offset), effective sensory uncertainty remains substantial for at least the visual modality (*σ*_*v*_ = 0.51, corresponding to a JND of approximately 200 ms). This result shows that visual duration discrimination itself was still imprecise despite no external visual noise. With these conditions, behavioral biases are close to linear functions of audiovisual conflict. As a result, these distinct integration strategies result in highly similar duration-discrimination patterns, making them difficult to distinguish based on behavior alone.

Consistent with this interpretation, the bimodal auditory duration judgments were less variable than predicted from the unimodal auditory and unimodal visual performance (Figure 17), and in some cases even less variable than the optimal-integration predictions based on unimodal noise estimates. We speculate that visual signals in the bimodal task contributed not only to duration information but also to interval segmentation. However, in unimodal auditory conditions, particularly under high noise, very short intervals may be difficult to detect, making it harder to segment the two intervals reliably, leading to increased response variability. Visual input may therefore stabilize auditory duration judgments beyond what would be expected from cue integration alone.

We emphasize that this lower-than-optimal variability is a separate phenomenon from the model comparison and does not bear on the choice among integration strategies. None of the candidate models anticipates lower-than-optimal bimodal variability; if anything, the cue-switching model predicts increased response variability because it relies on a single cue per trial. The reduced variability therefore points to an additional benefit of the visual signal—most plausibly more reliable interval segmentation—rather than to any particular integration rule.

### Causal inference versus heuristic strategies

In the present study, both the Bayesian causal-inference model and heuristic strategies were able to capture the main behavioral regularities, including the noise-dependent biases toward the visual duration. Notably, the probabilistic cue-switching model provided equal or superior fits to the behavioral data compared to the causal-inference model across multiple model-comparison criteria. At the same time, the causal-inference model did not produce uniquely diagnostic predictions within the experimentally tested conflict range. This convergence suggests that explicit inference about the causal structure does not contribute to the observed behavior.

Beyond providing the best overall fit, the cue-switching model offered an interpretable account of the noise dependence. The fitted probability of relying on the visual cue increased roughly threefold across noise conditions (Figure 12).

The lack of evidence for causal inference should not be interpreted as evidence against its plausibility. Rather, our results highlight a fundamental limitation in discriminating between integration strategies when sensory noise is sufficiently high. Under these conditions, the posterior probability of separate causes remains low even at large cue conflicts. As a result, bias toward the visual stimulus continues to increase approximately linearly with increasing cue conflict. In this regime, heuristic strategies that stochastically select between cues can closely approximate the predictions of more complex inferential models. Recent work has shown that probabilistic switching strategies can reproduce behavioral patterns often interpreted as Bayesian integration, even in the absence of explicit inference over causal structure (Ni and Ma, 2024). This perspective aligns with our finding that multiple integration strategies produce near-indistinguishable behavioral signatures in audiovisual-duration perception.

The difficulty of separating these strategies, however, does not undermine the preference for cue switching, because the two findings have different origins. Our model-recovery analysis shows that, in the empirically relevant parameter regime, the cue-switching model was highly identifiable (recovered in 84% of simulated datasets), whereas the forced-fusion and causal-inference models were mutually confusable. The limited separability we observe therefore reflects the near-equivalence of the two averaging-based models rather than a weakness of the cue-switching account. Consistent with this, cue switching was rarely selected when other models generated the data (in only 5 of 50 datasets generated by forced fusion and 5 of 50 generated by causal inference), so its empirical preference is a recoverable behavioral signature rather than an artifact of its additional flexibility.

### Asymmetry between shorter and longer audiovisual bias

Although most participants exhibited approximately linear bias patterns as a function of audiovisual conflict, the aggregated data revealed modest deviations from linearity at larger conflicts, particularly for conflicts in which visual duration exceeded auditory duration. These asymmetries were weaker or absent for conflicts in which the auditory duration was longer and did not produce the pronounced non-linear breakdown predicted by the classical causal-inference model.

The observed asymmetry in bias magnitude across opposing audiovisual conflicts was not anticipated by our initial hypotheses. Under a log-space encoding assumption, shorter durations are expected to be represented with lower uncertainty, which should lead to a stronger breakdown of integration for conflicts with a briefer stimulus. Contrary to this prediction, deviations from linearity were more apparent for large conflicts with a longer-duration visual stimulus.

One possible explanation is that factors beyond duration integration may influence performance at extreme conflicts. In particular, very short visual intervals may be less salient or less informative as temporal markers, reducing their influence on auditory judgments. Because this pattern was not predicted by the causal-inference model and was not consistently expressed at the individual-participant level, it should be interpreted cautiously. Rather than providing strong evidence for asymmetric causal inference, it highlights the sensitivity of duration judgments to stimulus interpretability and task structure.

## Conclusions

In this study, we investigated how auditory duration judgments are influenced by conflicting visual duration information under varying levels of sensory uncertainty. Across all experiments, we observed robust, noise-dependent biases toward visual duration, with bias magnitude increasing as auditory reliability decreased, consistent with reliability-weighted multisensory integration principles (Ernst and Banks, 2002; Van Wassenhove et al., 2008; Hartcher-O’Brien et al., 2014). These biases were largely linear across the tested conflict range and showed only modest asymmetries at larger conflicts. By comparing Bayesian causal-inference models with heuristic integration strategies, we found that no single computational mechanism was uniquely supported by the behavioral data: probabilistic cue-switching, causal inference, and forced-fusion models produced highly similar predictions within the experimentally plausible conflict range. This convergence reflects a fundamental limitation of model identifiability for audiovisual duration tasks: when encoding noise is high and behavioral biases remain approximately linear, distinct integration strategies yield near-indistinguishable behaviors. Importantly, these findings do not argue against the plausibility of causal inference in temporal perception, but instead suggest that short-duration audiovisual judgments may often be adequately explained by simpler heuristic strategies that approximate normative behavior without explicitly inferring causal structure (Caruso et al., 2018; Laquitaine and Gardner, 2018; Ni and Ma, 2024). More broadly, our results emphasize that understanding multisensory timing requires not only normative models but also careful consideration of task constraints, and the limits of model identifiability.

## Acknowledgments

This work was supported by NIH grant EY08266 and the NYUAD Center for Brain and Health, funded by Tamkeen under NYU Abu Dhabi Research Institute grant CG012.

## Appendix

### Model parameters

**Table 1.** Model parameter descriptions.

| Model | Lapse Rates | Sensory Noise | Prior $p_c$ | Other Params | Total $n$ |
| --- | --- | --- | --- | --- | --- |
| Forced fusion | 1 | 3 | – | – | 4 |
| Causal inference | 1 | 3 | ✓ | – | 5 |
| Probabilistic cue switching | 1 | 3 | – | $p_{v,l}, p_{v,h}$ | 6 |

*Note.* Sensory-noise parameters $\sigma_{a,l}$ , $\sigma_{a,h}$ , and $\sigma_v$ represent auditory (low and high noise) and visual standard deviations in log space. Prior parameter $p_c$ is the prior probability of common cause. Other parameters $p_{v,l}$ and $p_{v,h}$ are the switching-probability parameters in low and high auditory noise for the probabilistic cue-switching model.

### Task formulations

This section provides the mathematical formulations for the psychometric models used in each experimental task. We describe how duration estimates arise from noisy sensory measurements and how observers make comparison judgments.

#### Unimodal duration discrimination

In the unimodal tasks (auditory-only or visual-only), participants compared the durations of two sequentially presented stimuli: a standard stimulus with fixed duration (500 ms) and a test stimulus with variable duration determined by an adaptive staircase. The participant decides which interval was longer by comparing internal estimates of each duration.

##### Sensory encoding in log-space

To capture Weber-like scaling of temporal uncertainty (scalar variability), we assume that duration is encoded in logarithmic space and corrupted by additive Gaussian noise. Each duration measurement *m* is drawn from a Gaussian distribution centered on the log-transformed true stimulus duration:

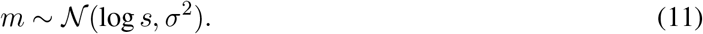

Under this encoding, measurements converted back to physical time are log-normally distributed, such that the standard deviation of perceived duration increases proportionally to the true duration.

#### Bayesian decoding (Gaussian prior case)

We describe the form of the posterior estimate under the assumption that log duration has a Gaussian prior, as this derivation underlies the precision-weighted combination used in all subsequent models. Let the measurement model be

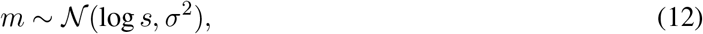

and assume a Gaussian prior over log-duration,

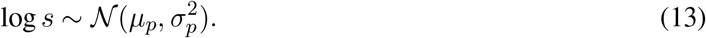

Under these assumptions, the posterior over log-duration is Gaussian. Throughout this paper we use *precision* (inverse variance) to simplify expressions involving reliability weighting. We write *J* = 1/*σ*^2^ for the sensory precision and 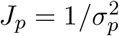 for the prior precision; more generally, 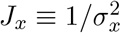 for any subscripted variance. The posterior mean (optimal estimator under squared-error loss in log-duration space) is then

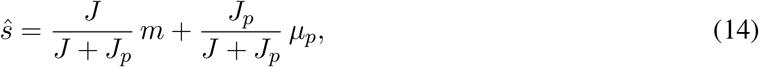

which can be written compactly as

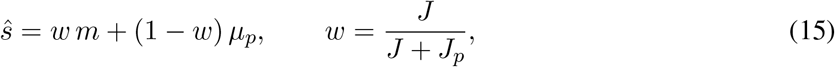

and the variance of the posterior is

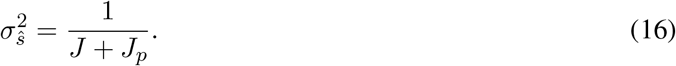

Here, all quantities 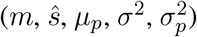 are in log-duration space.

For the standard and test intervals, the posterior means are

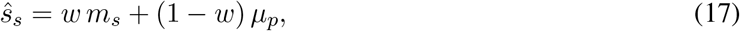

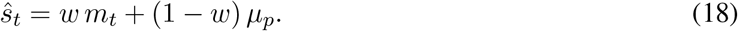

This formulation assumes that inference is performed in log-duration space, where sensory noise is Gaussian. For the Gaussian-prior example above, the posterior is Gaussian and its mean and mode coincide. The implemented models, however, use a bounded uniform prior and a BLS readout; consequently, the relevant estimate is the mean of the resulting truncated-normal posterior, as derived below.

#### Difference in posterior estimates

The difference between the two posterior estimates is therefore

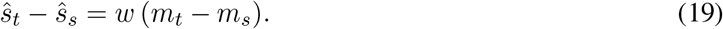

Because the same prior is applied to both intervals, the prior mean *µ*_*p*_ cancels out in the comparison. The prior thus acts symmetrically, compressing perceived differences toward zero by the factor *w <* 1, a form of regression toward the mean.

#### Decision rule

Decisions are based on the comparison of posterior estimates, such that the observer concludes that the test is longer when *ŝ*_*t*_ *> ŝ*_*s*_. Under the assumptions above, the posterior difference reduces to Eq.19, so that the decision is equivalent to comparing the measurements directly, i.e., determining whether *m*_*t*_ − *m*_*s*_ *>* 0. From the modeler’s perspective, the response probability is obtained by marginalizing over the measurement noise. Because *m*_*t*_ and *m*_*s*_ are independent random Gaussian variables with equal variance *σ*^2^, their difference follows

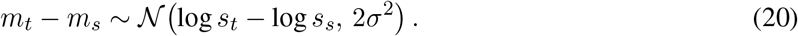

Therefore,

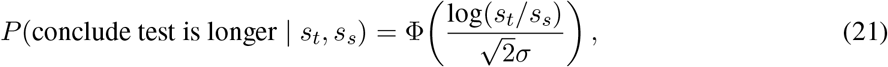

where Φ(·) denotes the cumulative distribution function of the standard normal distribution.

#### Psychometric function

Because test and standard are presented in random order, the psychometric function cannot have observer bias. However, we do allow for occasional lapses, including a lapse rate *λ* to account for them. The psychometric function used for the unimodal fits is thus:

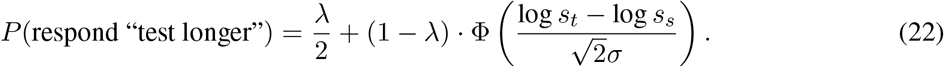

#### Cross-modal duration comparison

To quantify modality-specific biases in duration perception, participants completed a cross-modal duration-comparison task in which they judged whether an auditory or visual interval appeared longer. Responses were fit with cumulative log-normal psychometric functions using maximum-likelihood estimation.

The point of subjective equality (PSE; parameter *µ*) quantified the modality-specific duration bias, corresponding to the physical offset required for the two modalities to be perceived as equal in duration. The slope parameter quantified discrimination sensitivity. Individual PSE estimates were subsequently used to calibrate the visual durations in the main bimodal experiment, ensuring that the nominal zero-conflict condition corresponded to perceptually matched auditory and visual durations.

### Models of audiovisual duration estimation

#### Forced fusion

Our forced-fusion model assumes that participants estimate auditory log duration by integrating visual and auditory measurements in log-duration space, weighted by their relative reliabilities. Before applying the bounded uniform prior, the product of the auditory and visual Gaussian likelihoods is proportional to a Gaussian density with mean and standard deviation

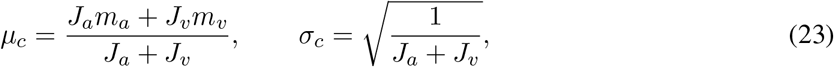

where *m*_*a*_ and *m*_*v*_ denote noisy sensory measurements in log-duration space, and 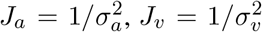 are the corresponding precisions (with *σ*_*a*_ ∈ {*σ*_*a,l*_, *σ*_*a,h*_} depending on the condition).

The implemented readout applies the same bounded uniform prior over log-duration, [*t*_min_, *t*_max_], used in the causal-inference model. The resulting posterior is a normal distribution with parameters *µ*_*c*_ and *σ*_*c*_, truncated to these bounds. The forced-fusion estimate is its Bayes least-squares (BLS) estimate, i.e., its posterior mean:

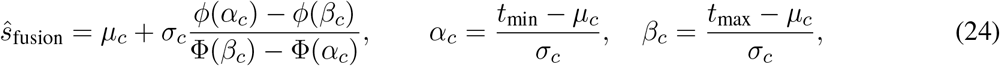

where *ϕ* and Φ are the standard normal density and cumulative distribution functions. Thus, the implemented estimate is a posterior mean rather than a maximum a posteriori (MAP) estimate. Away from the prior bounds, the truncation correction becomes negligible and the estimate approaches the usual reliability-weighted mean *µ*_*c*_.

Here the forced-fusion model serves as an important benchmark: it predicts that PSE shifts should follow a linear relationship with audiovisual conflict, with slope determined solely by cue reliability. This represents the limiting case where the observer *always* assumes a common cause, regardless of sensory evidence. In forced-fusion models, this estimate is used on every trial, leading to a linear relationship between audiovisual conflict and bias. In contrast, Bayesian causal-inference models combine this fused estimate with segregated estimates according to the posterior probability of a common cause, *p*(*C* = 1 | *m*_*a*_, *m*_*v*_). Because this posterior probability decreases with increasing conflict, the effective integration strength becomes conflict-dependent, producing nonlinear bias functions. Before truncation, the common-cause posterior variance is 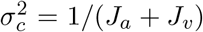, which is smaller than either measurement variance. The bounded prior further restricts the posterior support.

#### Probabilistic cue switching

The probabilistic cue-switching model represents an alternative strategy. On each trial, the observer stochastically selects *one* modality and bases the estimate entirely on that cue. For the selected modality, the implementation uses the BLS estimate under the same bounded uniform prior: the mean of the single-cue truncated-normal posterior. Thus, the selected cue is decoded using a posterior mean rather than its raw measurement or a MAP estimate. This contrasts with averaging-based strategies that combine information from both modalities. The probability of using the visual cue is a free parameter of the model.

#### Comparison between switching and integration

The key behavioral distinction between switching and causal inference lies in the distribution of responses across repeated trials at a fixed conflict level. Switching generates different estimate distributions than the fusion and causal-inference models. Under switching, the decision on each trial is based on the posterior-mean estimate derived from either the auditory or the visual measurement. Across trials, this produces a mixture of decisions. From the modeler’s perspective, this mixture results in psychometric functions that can resemble reliability-weighted integration when averaged across trials, even though no cue averaging occurs within a single trial. Averaging models imply a single fused internal estimate on each trial, centered near the reliability-weighted combination of the true stimulus durations. In contrast, switching models imply cue selection on each trial.

### Bayesian causal-inference model

#### Likelihood of a common cause

If the model assumes that both durations arise from a common source, there is a single latent log-duration *y* = log *s*, and the measurements (in log-duration space) are:

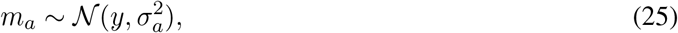

and

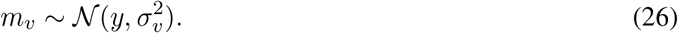

Let *s*_min_ and *s*_max_ denote the physical-duration bounds fixed from the range of test durations, and define the corresponding log-space bounds as *t*_min_ = log *s*_min_ and *t*_max_ = log *s*_max_. We assume that the participant uses a flat prior over the latent log-duration *y* on [*t*_min_, *t*_max_]. This prior is needed to normalize the integral below to compute the evidence for a common cause. The prior is

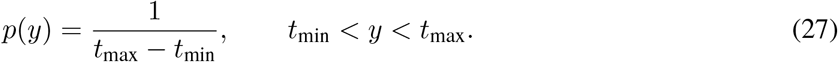

To compute the evidence for a common cause, we marginalize over the unknown latent duration *y*:

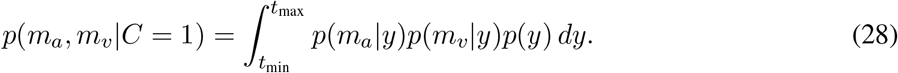

Let Δ_*t*_ = *t*_max_ − *t*_min_. Taking the prior out of the integral:

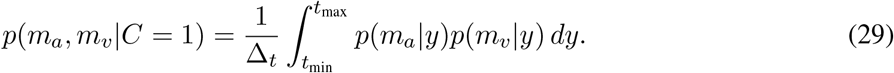

Writing the likelihoods as Gaussians:

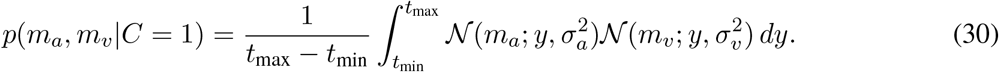

Because a Gaussian is symmetric in its arguments:

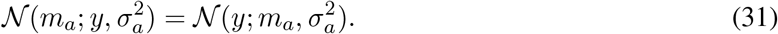

It is identical to write the likelihood as:

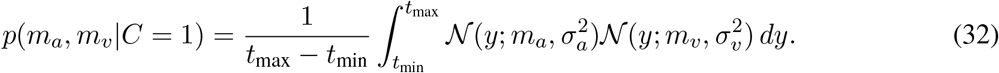

#### Product of two Gaussians and the final likelihood

The integrand is the product of two Gaussians in the same variable *y*. By completing the square in *y* (using the precisions *J*_*a*_ and *J*_*v*_ defined earlier), this product factors into a Gaussian in *y* times a term that is constant with respect to *y*:

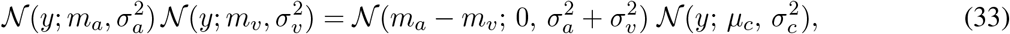

where the precision-weighted mean and variance of the fused posterior are

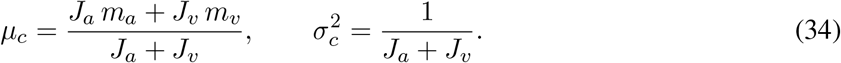

The first factor, 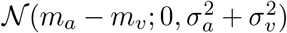, is a constant with respect to *y* that quantifies the compatibility of the two measurements. Substituting this factorization into the integral and evaluating the Gaussian over the finite prior support gives:

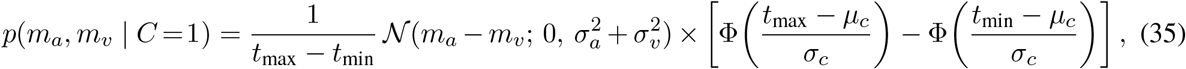

where Φ is the standard normal cumulative distribution function.

#### Separate causes

Under the separate-causes hypothesis, we estimate two latent durations, one for each modality (*y*_*a*_ and *y*_*v*_).

These latent durations *y*_*a*_ and *y*_*v*_ are independent of each other but both are bounded by the same *t*_min_ and *t*_max_, and they generate measurements from noisy Gaussian distributions:

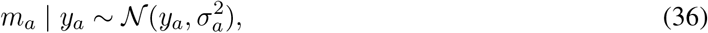

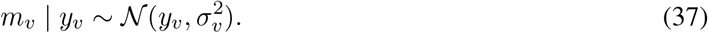

Because they are bounded by the same minimum and maximum durations, the uniform prior densities are:

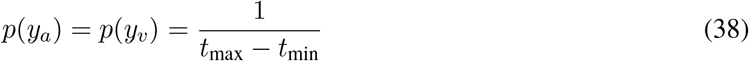

over the range [*t*_min_, *t*_max_].

The likelihood of the two measurements under *C* = 2 is:

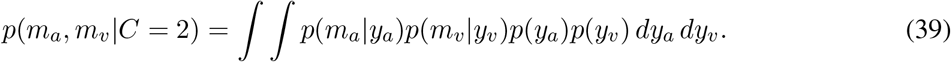

#### Factor the double integral

Because the integrand separates into a function of *y*_*a*_ times a function of *y*_*v*_ and the domains (boundaries) are independent:

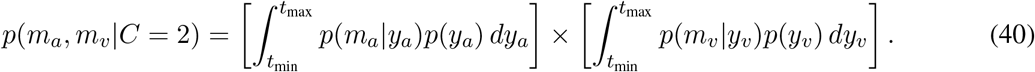

Because the integrands are Gaussians in the latent variables, each integral evaluates to a Φ-difference as in the common-cause case:

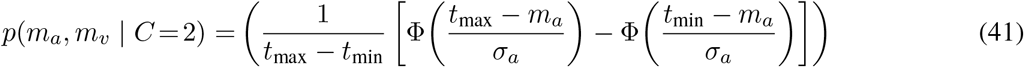

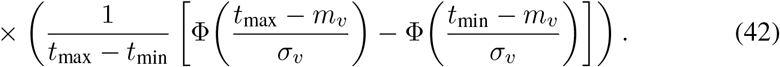

#### Posterior probability of common cause

The posterior probability of a common cause is obtained by combining the common-cause and separate-causes evidences with the prior probability of common cause:

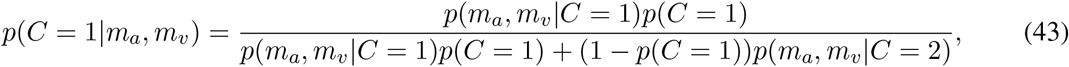

where *p*(*C* = 1) is the prior probability of common cause.

#### Estimate with causal inference

The bounded uniform prior above is used both to compute the marginal likelihoods and to decode duration under each causal structure. The duration readout is the Bayes least-squares estimate (posterior mean) under squared-error loss. Under a common cause, first define the precision-weighted Gaussian parameters

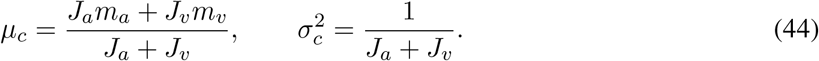

After applying the bounded uniform prior, the posterior is this Gaussian truncated to [*t*_min_, *t*_max_]. Its mean is

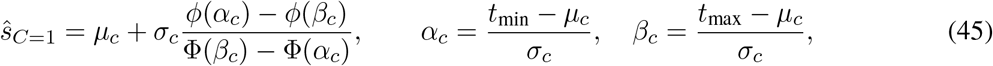

where *ϕ* and Φ are the standard normal density and cumulative distribution functions, respectively. Under separate causes, the task-relevant auditory posterior is a Gaussian centered on *m*_*a*_, truncated to the same bounds, and its mean is

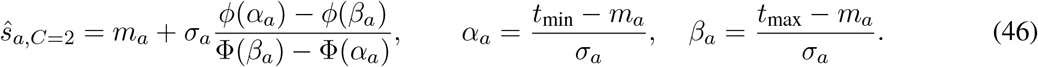

The final auditory-duration estimate is the posterior-probability-weighted average of these two BLS estimates:

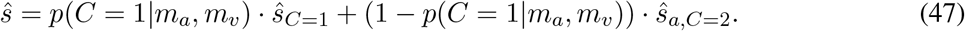

Here, 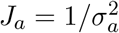 and 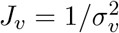 denote sensory precisions (with *σ*_*a*_ ∈ {*σ*_*a,l*_, *σ*_*a,h*_} depending on condition). All measurements, bounds, and estimates in these expressions are in log-duration space. In the segregated case (*C* = 2), only the auditory posterior contributes to the task response, consistent with the instruction to judge auditory duration.

### Model fits to individual subjects’ data

Figure 14 shows point of subjective equality (PSE) shifts as a function of audiovisual duration conflict for individual participants, plotted separately for the low and high auditory-noise conditions. This figure illustrates individual differences in bias magnitude and noise dependence, as well as the substantial overlap in predicted PSE–conflict functions across models within the tested conflict range.

**Figure 14.**
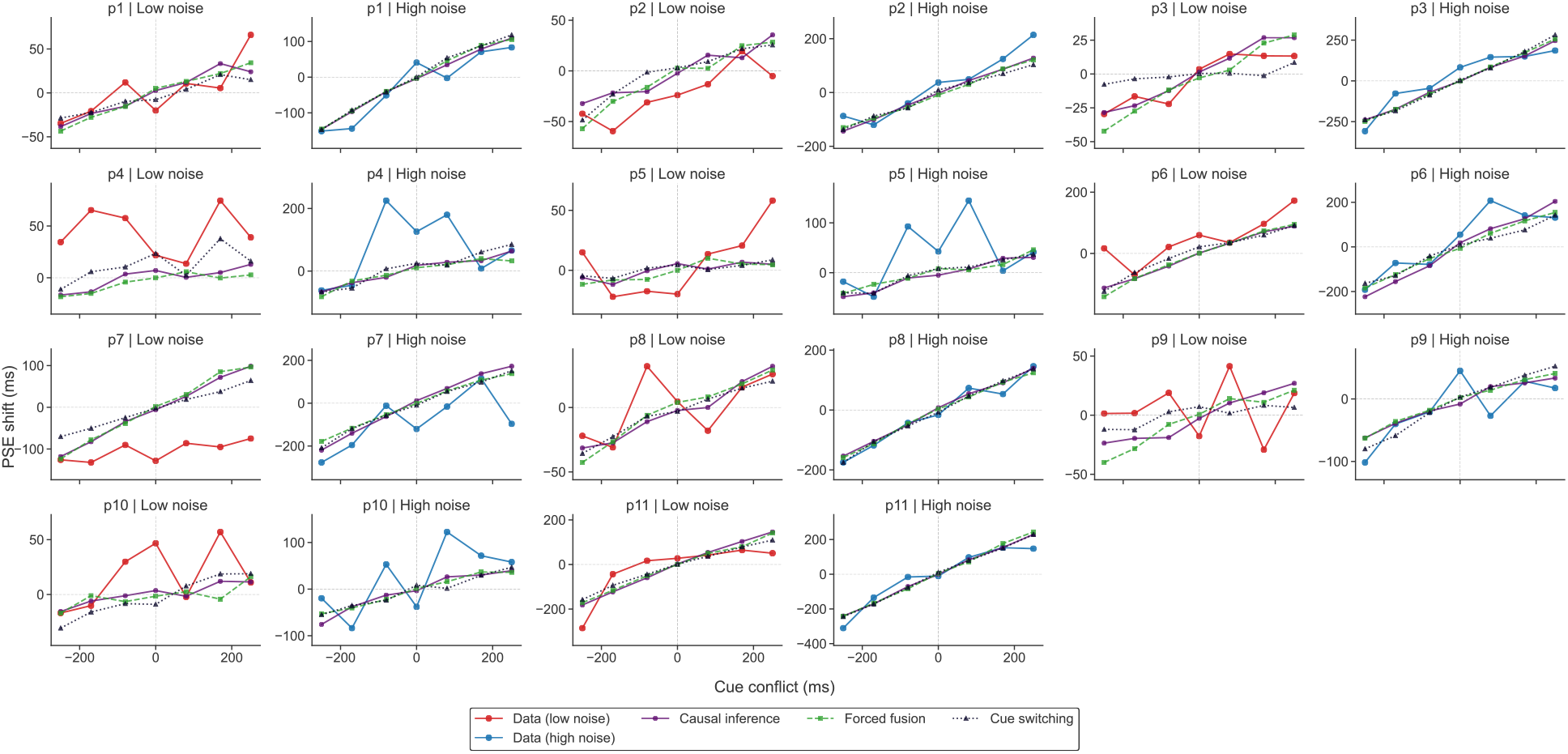
Measured and predicted PSE shifts as a function of conflict levels across different auditory-noise conditions for each participant. Here, the *σ* values are estimated using the bimodal data.

**Figure 15.**
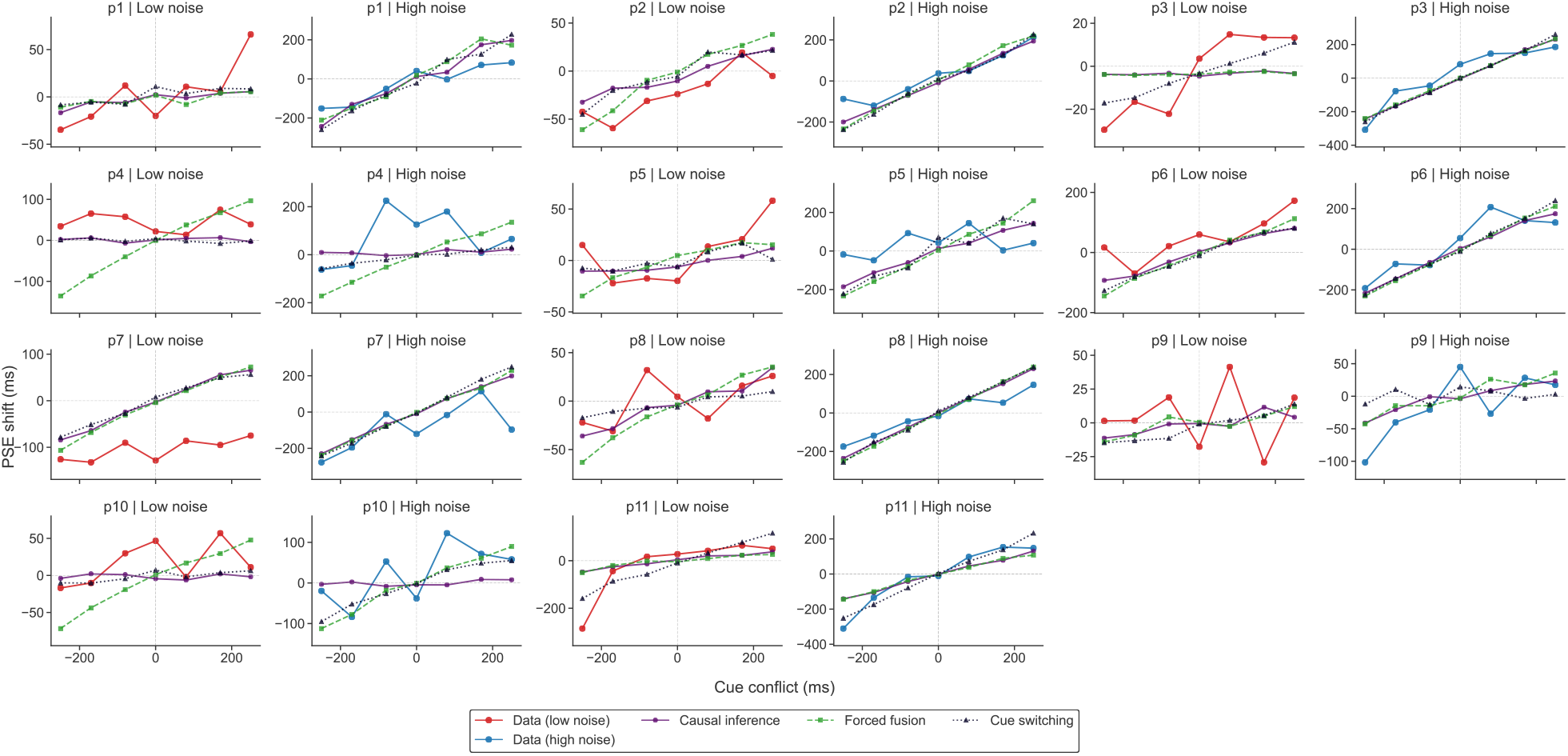
Same as Figure 14, except that here the models are fitted using the sensory-noise parameters (*σ*_*a,l*_, *σ*_*a,h*_, *σ*_*v*_) that were estimated based on the psychometric fits to the unimodal data.

### Posteriors and likelihoods of common causes

To further illustrate how the causal-inference model interprets audiovisual conflict at the individual-participant level, we plot the log-likelihoods of the common-cause and separate-causes hypotheses and the resulting posterior probability of a common cause as a function of conflict magnitude under the fitted log-space causal-inference model (Figure 16). Across participants, the likelihood of a common cause remained comparable to or higher than that of separate causes, so the posterior probability of a common cause stayed relatively high across the tested conflict range, particularly in the high-noise condition. All quantities were computed via Monte Carlo simulation using 2,000 samples per conflict level and participant-specific sensory-noise estimates (*σ*_*a*_, *σ*_*v*_) and common-cause prior (*p*_*c*_) from the log-normal causal-inference model fit. Across participants,the posterior remains close to the prior for nearly all observers, indicating that the fitted parameters leave the causal-inference mechanism unable to distinguish common from separate causes at the conflict levels tested.

**Figure 16.**
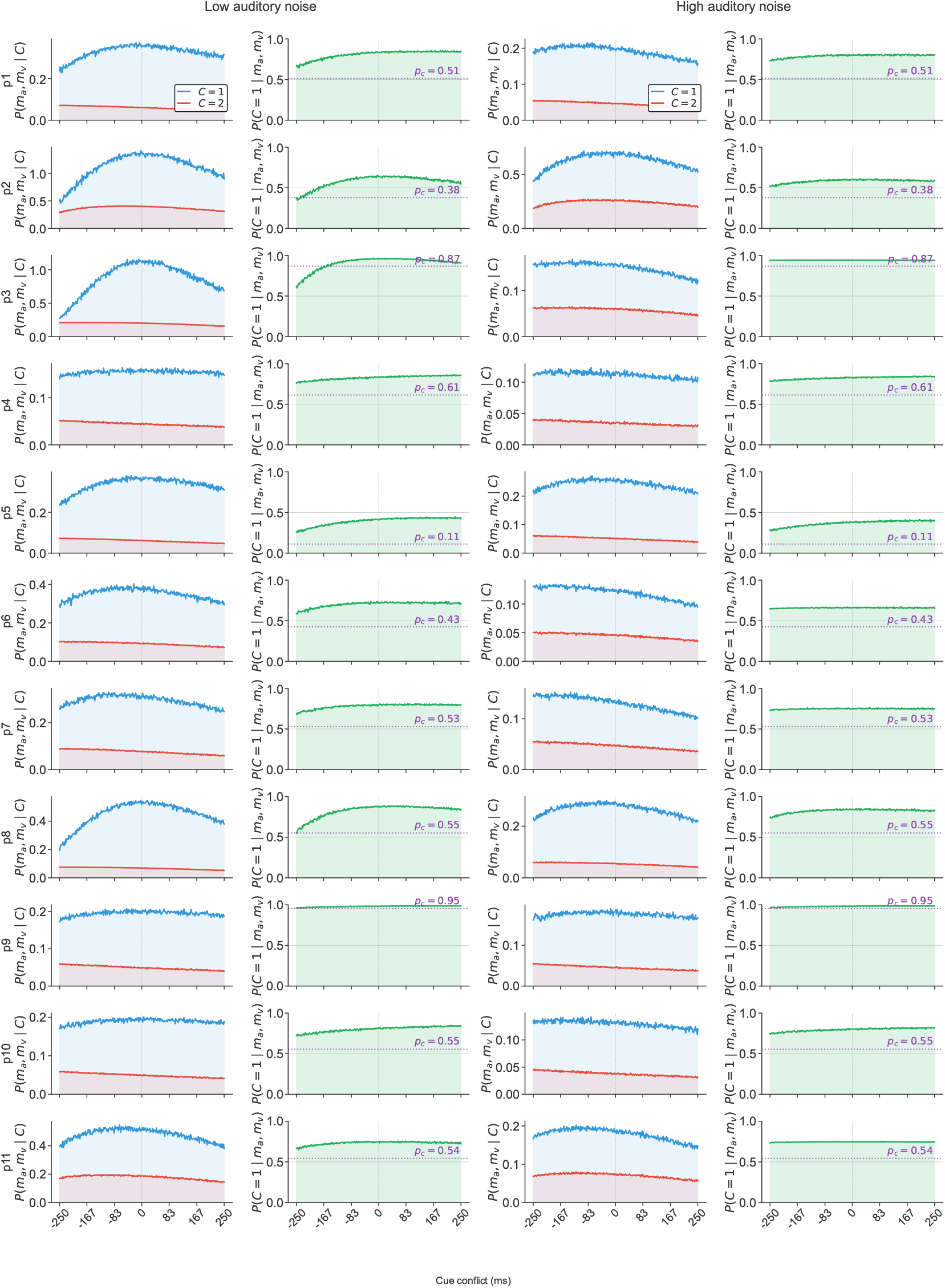
Likelihoods and posterior probability of a common cause for each participant. Left two columns: low auditory noise; right two columns: high auditory noise. Columns 1 and 3: the likelihood of the measurements under a common cause, *P* (*m*_*a*_, *m*_*v*_ | *C*=1) (blue), and under separate causes, *P* (*m*_*a*_, *m*_*v*_ | *C*=2) (red), are plotted on the same axes. Columns 2 and 4: the posterior probability of a common cause, *P* (*C*=1 | *m*_*a*_, *m*_*v*_), with the dotted line indicating the fitted prior *p*_*c*_. Each row corresponds to one participant.

### Unimodal-derived predictions of bimodal uncertainty

Figure 17 compares empirically observed audiovisual variability (*σ*_*AV*_) with variability predicted under optimal reliability-weighted integration (*σ*_opt_), derived from unimodal sensory-noise estimates. This calculation is used only for the comparison of observed and predicted uncertainty.

**Figure 17.**
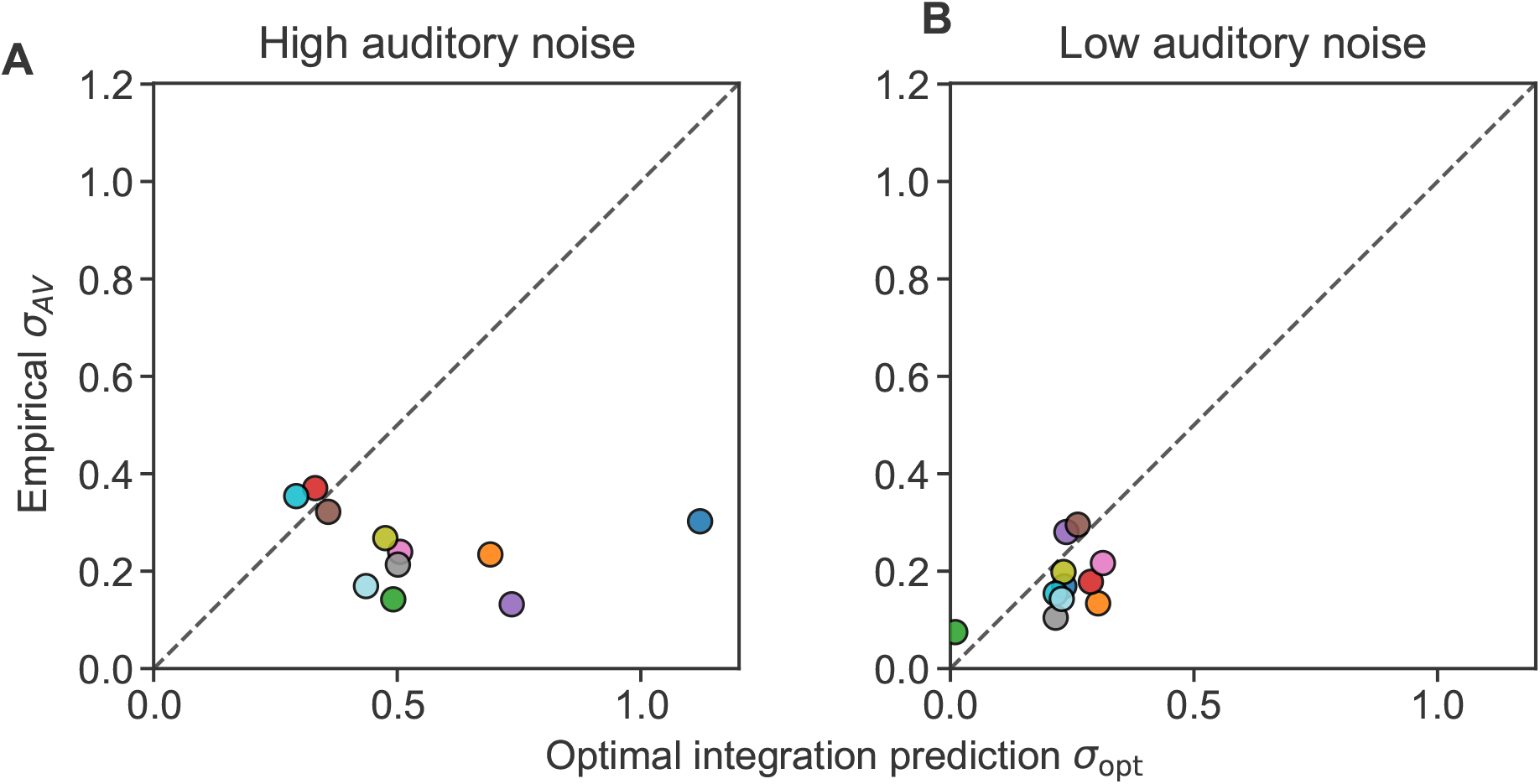
Comparison of predicted optimal-integration variability with observed audiovisual variability. Each panel compares empirically observed audiovisual variability (*σ*_*AV*_ ; obtained by fitting psychometric functions to the bimodal data, *y*-axis) with variability predicted by optimal reliability-weighted integration (*σ*_opt_, *x*-axis), computed from unimodal psychometric-fit noise estimates (*σ*_*a,h*_ and *σ*_*a,l*_ for auditory high- and low-noise conditions, and *σ*_*v*_ for vision). Each point is one participant; the dashed line indicates equality (*σ*_opt_ = *σ*_*AV*_).

Under the assumption of statistically optimal cue combination with independent Gaussian measurement noise and no additional duration-prior variance term, the predicted audiovisual variance is given by

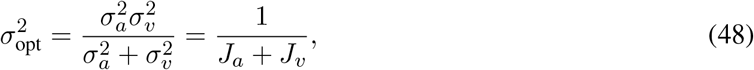

where *σ*_*a*_ and *σ*_*v*_ denote auditory and visual noise, respectively, and 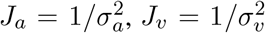 are the corresponding precisions.

Observed audiovisual variability (*σ*_*AV*_) was estimated by fitting psychometric functions to the bimodal data. Predictions are computed separately for the high- and low-noise auditory conditions using the corresponding unimodal estimates (*σ*_*a,h*_ and *σ*_*a,l*_), and compared to the fitted bimodal variability.

Panels show the results for each noise condition. Each point corresponds to an individual participant. The dashed unity line (*σ*_opt_ = *σ*_*AV*_) indicates perfect agreement between predicted and observed variability. Systematic deviations from this line reflect departures from optimal integration based solely on unimodal noise estimates.

Interestingly, observed audiovisual noise was substantially lower than the noise predicted by optimal reliability-weighted integration based on unimodal sensory estimates, particularly in the high auditory-noise condition. In several participants, bimodal uncertainty fell well below the unity line, indicating higher precision than predicted from unimodal noise alone.

One possible explanation is that the visual stimulus in the bimodal task provided highly precise temporal anchors marking stimulus onset and offset. These visual transients may reduce temporal uncertainty beyond simple duration integration by improving the detection of the interval itself. As a result, auditory duration judgments in the bimodal task may benefit from additional temporal structure (precise cue for onset and offset of the event) that is absent in the unimodal auditory condition, leading to lower overall variability than predicted from unimodal noise estimates alone.

A second possibility is that sensory precision should not be expected to stay stationary across experiment and trials. The bimodal task involved substantially more trials and longer exposure to the stimulus structure, potentially leading to implicit training on temporal discrimination over time. If unimodal noise estimates reflect earlier performance, they may overestimate the effective sensory variability present during the bimodal experiment.

These findings suggest that unimodal-derived sensory noise parameters may not fully generalize to multisensory contexts, particularly when additional temporal cues are available.

### Model recovery

Model recovery is a crucial validation step that tests whether our model-comparison procedure can correctly identify the true data-generating model. This ensures that when we select a “best model” for real data, we’re doing so reliably. To perform this analysis, after fitting each candidate model to the empirical data, we used the fitted parameter values to generate 50 synthetic datasets per model. Parameters were sampled from group-level distributions estimated from the 11 participants included in the main analyses. For each sampled parameter set, we simulated trial-by-trial responses according to the probabilities predicted by the generating model’s psychometric function. Each synthetic dataset matched the original experimental design (same stimulus conditions, trial counts, and structure). We then fit all candidate models to every simulated dataset and recorded their AIC values and the winning model.

The model-recovery results reveal substantial differences in identifiability across models (Figure 18). Recovery was strongest for the probabilistic cue-switching strategy, which was correctly identified in 84% of simulated datasets, indicating that this model generates relatively distinctive behavioral predictions. Recovery for the forced-fusion model was moderate (74%), with appreciable confusion with the causal-inference model. In contrast, the causal-inference model showed poor mutual discriminability; 44% of the time the recovered model was correct and also 46% of the time the results were confused with the forced-fusion model. This pattern suggests that only the probabilistic cue-switching model generates distinguishable behavioral data.

**Figure 18.**
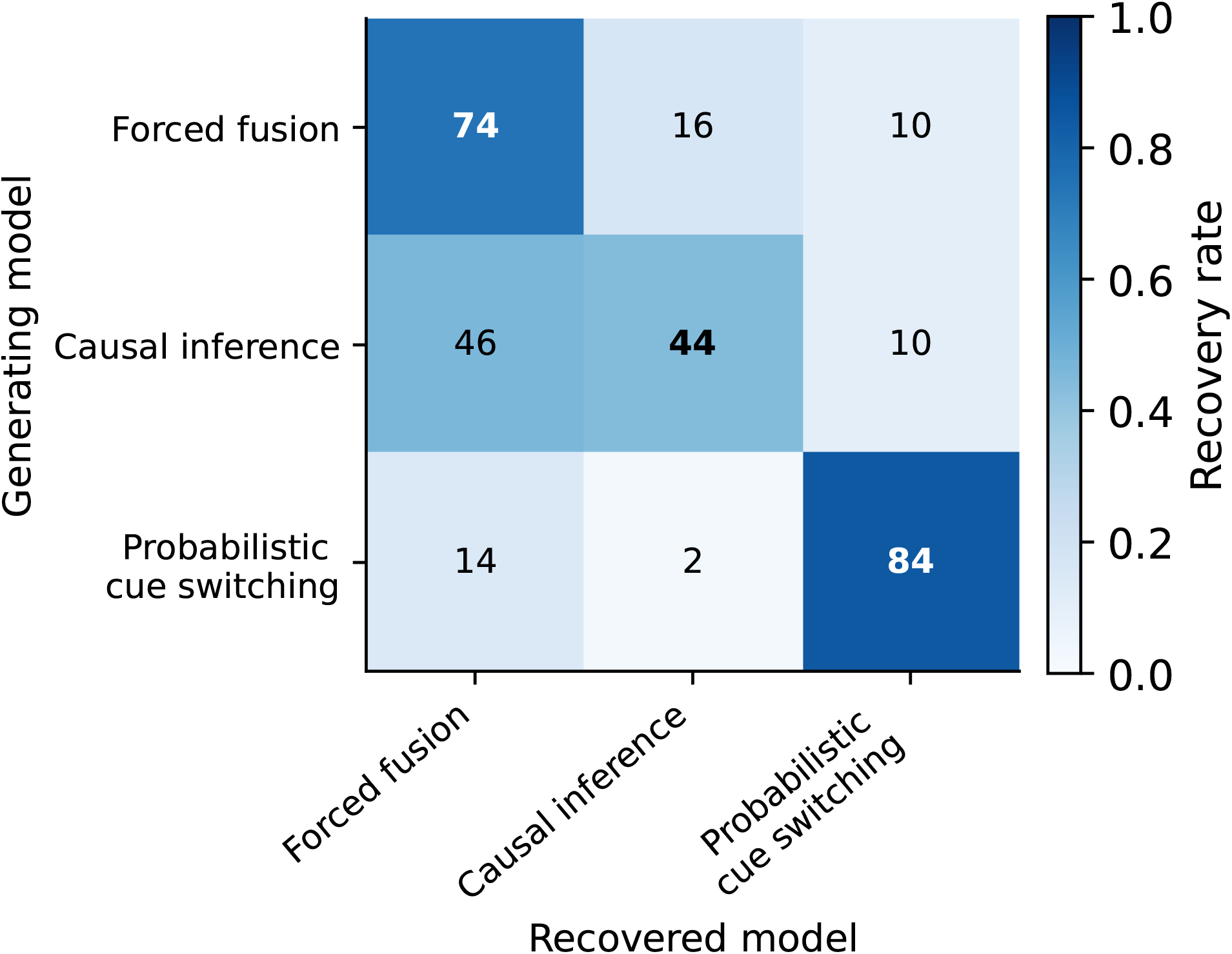
Model recovery. Validation of model identifiability through synthetic data analysis. For each of the three candidate models (rows), we generated 50 synthetic datasets using parameters sampled from the group-level distribution (mean *±* SD from 11 participants). All models were then fit to each synthetic dataset, and the best-fitting model was determined by AIC. Row-normalized recovery rates show the percentage of times when a dataset generated by one model was identified as coming from each of the three models. Values on the diagonal indicate successful recovery of the true generating model. The diagonal values reveal important differences in model identifiability: probabilistic cue switching showed very good recovery, while averaging-based models (forced fusion, causal inference) showed weaker recovery, indicating that these models make similar predictions and are less distinguishable using our behavioral data.

Importantly, the high recovery of the cue-switching model did not reflect a general tendency to over-select the most flexible model. Cue switching was selected for only 5 of 50 datasets generated by the forced-fusion model and 5 of 50 datasets generated by the causal-inference model, but it was selected for 42 of 50 datasets generated by the cue-switching model. Its preference in the empirical data therefore reflects a genuine, recoverable behavioral signature rather than spurious model flexibility.

### Parameter recovery

Using the same synthetic datasets, we also conducted a parameter-recovery analysis to ask whether parameters could be recovered when the generating model class was known. The parameter-recovery results (Figure 19) show a graded pattern rather than uniformly high recovery across all parameters. Across models, the lapse rate (*λ*) and the low-auditory-noise parameter (*σ*_*a,l*_) were recovered accurately, with high correlations between true and recovered values and slopes close to the identity line. Most importantly for the cue-switching account, the two switching-probability parameters were recovered extremely well (*p*_*v,l*_: *r* = 0.98; *p*_*v,h*_: *r* = 0.92), indicating that the model’s estimates of noise-dependent reliance on the visual duration are strongly constrained by the behavioral data.

**Figure 19.**
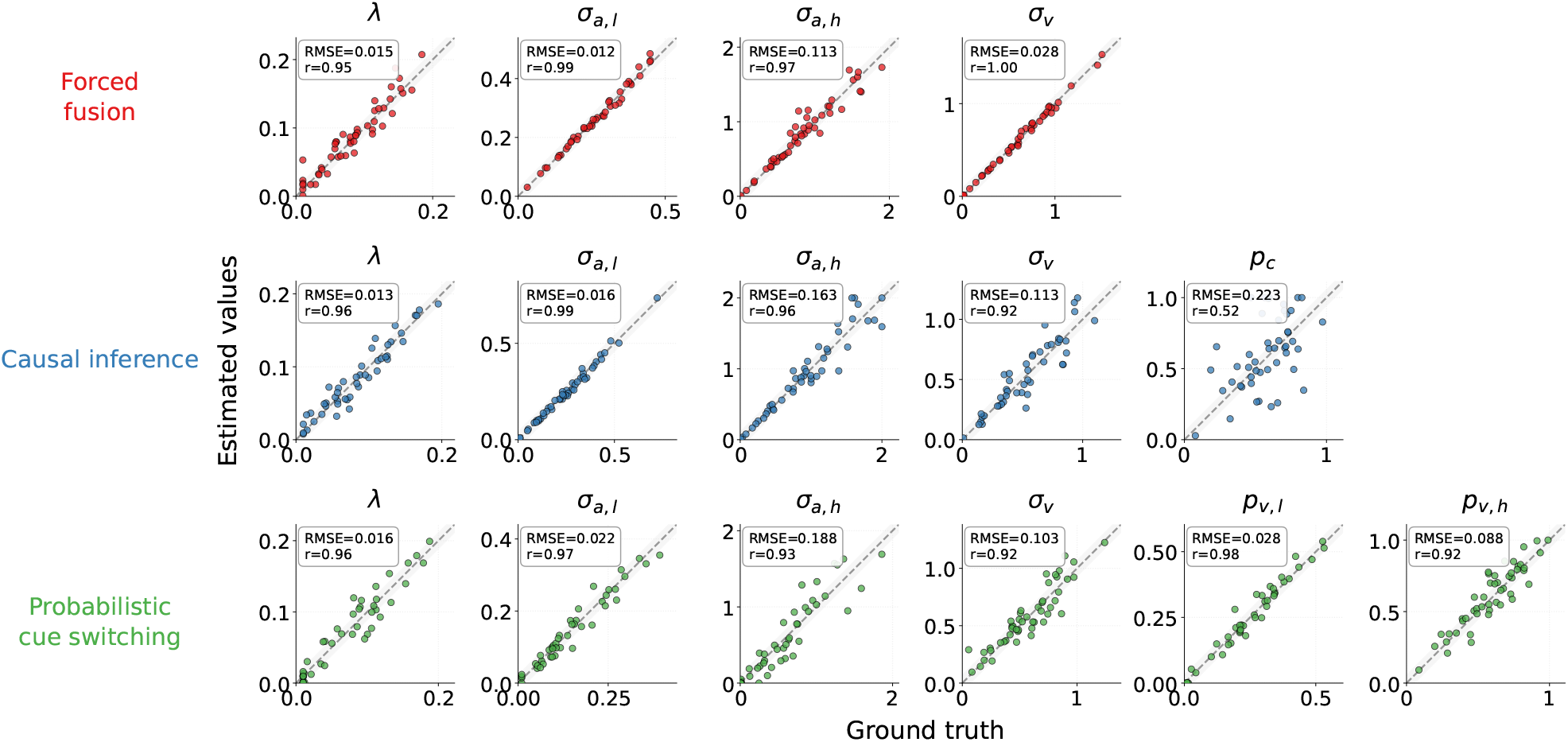
Parameter-recovery analysis for model validation. Each row corresponds to a generative model and each column to a model parameter. Synthetic datasets (*N* = 50) were generated from fitted group-level parameter distributions and refit with the same model. Scatterplots show true versus recovered parameter values; dashed diagonals indicate perfect recovery and gray bands show a *±* 5% tolerance region. Insets report Pearson correlation (*r*) and RMSE. Recovery was strongest for the lapse rate, low-auditory-noise parameter, and cue-switching probabilities; high-auditory-noise and visual-noise parameters were recovered less precisely, and the causal-prior parameter showed weak recovery.

**Figure 20.**
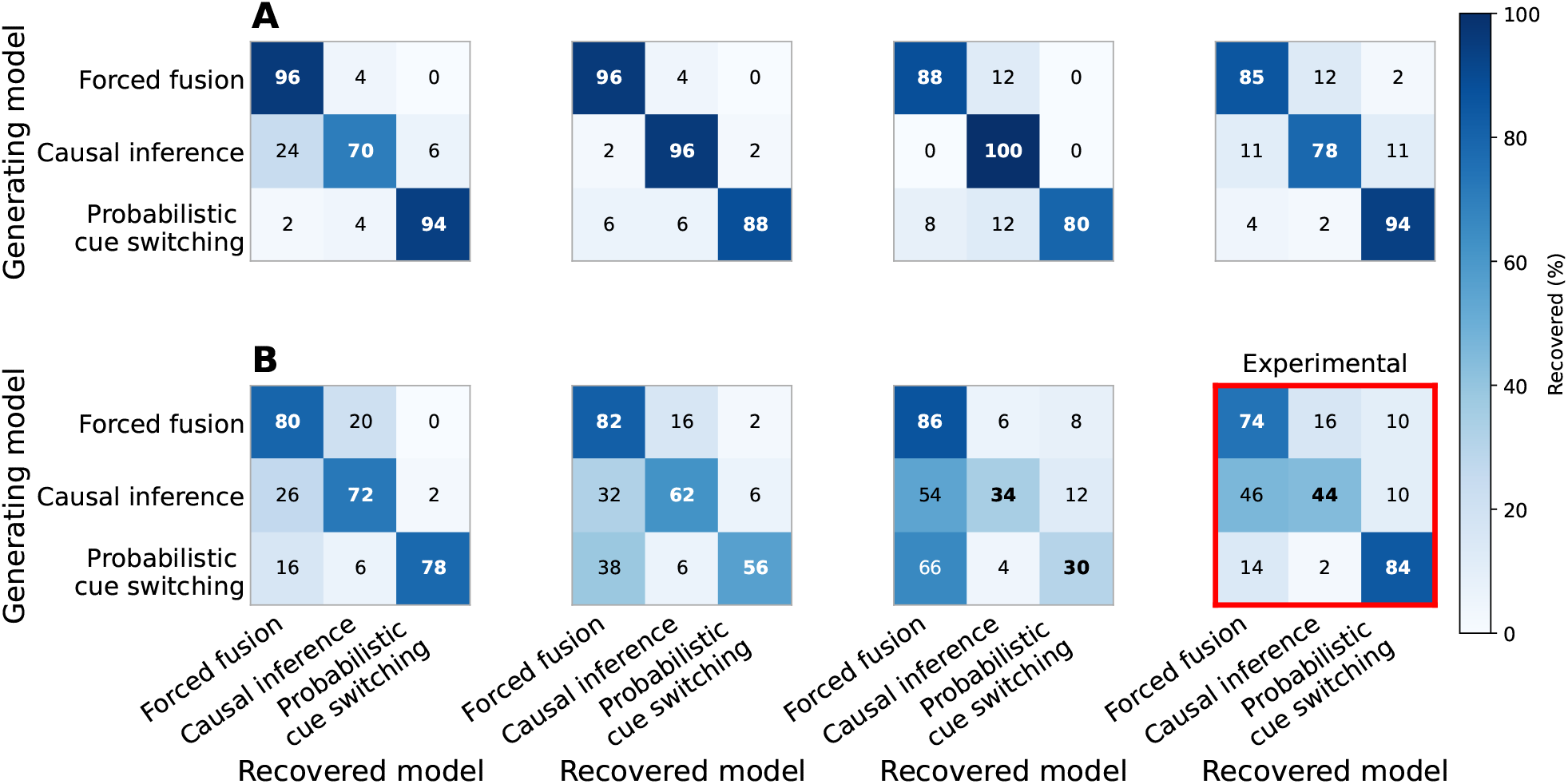
Model recovery across sensory-noise and conflict regimes. Confusion matrices for the three candidate models: forced fusion, causal inference, and probabilistic cue switching. Rows indicate the generating model, and columns indicate the recovered model selected by AIC. Cell values show row-normalized recovery percentages, with diagonal entries corresponding to correct recovery. Row **A** (top) uses conflict_max_ = 0.45 s, whereas row **B** (bottom) uses conflict_max_ = 0.25 s. The first three columns use uniformly sampled synthetic sensory-noise regimes: *σ*_*a*_, *σ*_*v*_ ∼ *U* [0.01, 0.20], *U* [0.20, 0.40], and *U* [0.30, 0.70]. The rightmost column is empirically informed but uses different sampling procedures across rows. The lower-right panel, highlighted in red, uses parameters sampled from the fitted group-level distributions and reproduces the experimentally tested conflict range. The upper-right panel uses uniform sampling over the empirical fitted ranges (*σ*_*a*_ ∈ [0.12, 0.66], *σ*_*v*_ ∈ [0.26, 1.13]) with the wider conflict range. Consequently, the rightmost column lies off the synthetic noise gradient and its two panels should not be interpreted as identically sampled. Recovery generally decreased with increasing sensory noise and narrower conflict ranges. In the lower-right empirical recovery panel, probabilistic cue switching showed strong recovery, whereas forced fusion and causal inference exhibited greater mutual confusability.

In contrast, recovery was less precise especially for the visual-noise parameter (*σ*_*v*_) in the causal-inference and cue-switching models. This weaker recovery of *σ*_*v*_ likely reflects the structure of the task and model: participants judged the auditory interval, so visual reliability influenced responses only indirectly through conflict (difference) in auditory and visual duration. As a result, changes in *σ*_*v*_ can be partially absorbed by other parameters, and the fitted value of *σ*_*v*_ can trade off with parameters controlling visual influence, such as *σ*_*a*_, *p*_*c*_, or *p*_*v*_. The causal-prior parameter (*p*_*c*_) showed low recovery, indicating that individual estimates of this parameter should be interpreted cautiously.

Thus, parameter recovery supports the reliability of the main response-shaping components of the models, especially auditory precision, and the cue-switching probabilities, but it does not imply that every fitted parameter is equally identifiable. This pattern refines the interpretation of the model-recovery results. The difficulty of distinguishing forced fusion from causal inference likely reflects both the overlap in their behavioral predictions within the tested conflict range and the weak recovery of parameters, such as *p*_*c*_ and *σ*_*v*_, that would otherwise help separate causal-inference behavior from averaging-like behavior. By contrast, the excellent recovery of *p*_*v,l*_ and *p*_*v,h*_ supports the interpretation that the empirical preference for probabilistic cue switching reflects a recoverable behavioral signature rather than an artifact of poorly estimated switching parameters.

### Extended simulations across sensory-noise and conflict regimes

To assess when the candidate models should be expected to become distinguishable, and under which conditions the hypotheses about cue integration could be disentangled, we performed an extended model-recovery analysis across simulated sensory-noise and conflict sampling regimes. We simulated datasets from the three candidate models while varying both the sensory-noise range and the maximum audiovisual conflict. Three synthetic sensory-noise regimes were sampled uniformly: very low noise, *σ*_*a*_, *σ*_*v*_ ∼ *U* [0.01, 0.20]; low noise, *σ*_*a*_, *σ*_*v*_ ∼ *U* [0.20, 0.40]; and medium noise, *σ*_*a*_, *σ*_*v*_ ∼ *U* [0.30, 0.70]. The rightmost column contains two empirically informed analyses that used different sampling procedures. For the experimentally tested maximum conflict (0.25 s), parameters were sampled from the fitted group-level distributions. For the wider maximum conflict (0.45 s), sensory-noise parameters were sampled uniformly over the empirical fitted ranges (*σ*_*a*_ ∈ [0.12, 0.66] and *σ*_*v*_ ∈ [0.26, 1.13]).

The grid recovery analysis should be read as a map of model identifiability across possible experimental regimes. Each panel shows how often the generating model was recovered after refitting simulated datasets with all three candidate models. Higher diagonal values indicate better model recovery, whereas off-diagonal values indicate systematic confusions between models. Sensory uncertainty increases from left to right across the first three uniformly sampled synthetic columns. The empirically informed rightmost column is not a continuation of this synthetic-noise axis, and its two panels should not be treated as identically sampled. Moving from the top row to the bottom row reduces the maximum conflict range from 0.45 s to 0.25 s.

This analysis shows that model discriminability depends strongly on both sensory noise and the tested range of audiovisual conflicts. Recovery was generally best when sensory noise was low and the conflict range was wide, and poorest when sensory noise was higher and the conflict range was narrow. Importantly, the lower-right empirical recovery panel shows that the experimentally observed regime was one in which forced fusion and causal inference were difficult to distinguish, whereas probabilistic cue switching remained relatively identifiable.

When reading the cue-switching row across columns, it is important not to treat all four panels as a single monotonic noise sweep. Across the three synthetic regimes, recovery of the cue-switching model declined steadily as sensory noise increased (for the 0.25 s conflict range, from 86% in the very-low-noise regime to 48% and 28% in the low- and medium-noise regimes), with its simulated datasets increasingly misclassified as forced fusion. The lower-right empirical recovery panel does not continue this downward trend: its high recovery (84%) reflects parameters sampled from the fitted group-level distributions, under which cue switching produces a particularly distinctive behavioral signature. Its placement next to the uniformly sampled synthetic regimes should therefore not be interpreted as a non-monotonic dependence on sensory noise.

Model recovery generally improved as sensory noise decreased and as the maximum conflict range increased from 0.25 s to 0.45 s. This pattern suggests that larger audiovisual discrepancies and lower sensory uncertainty provide more diagnostic information for distinguishing between integration strategies. For the experimentally tested conflict range, where the maximum conflict was 0.25 s, recovery remained more limited, especially for distinguishing forced fusion from causal inference. However, probabilistic cue switching remained comparatively identifiable in the empirical recovery analysis.

One exception to the general trend occurred under the wider 0.45 s conflict range, where recovery in the very-low-noise regime was slightly poorer than in the low-noise regime. This reduction was driven partly by an asymmetric confusion: 24% of datasets generated by the causal-inference model were classified as forced fusion. Inspection of the fitted parameters revealed a compensatory mechanism underlying this confusion (Figure 21). When forced fusion was fitted to the 50 causal-inference-generated datasets, its estimate of visual sensory noise exceeded the generating value in every dataset. Median *σ*_*v*_ increased from 0.085 in the generating model to 1.085 in the forced-fusion fit, corresponding to a median paired increase of 1.008 (bootstrap 95% CI [0.825, 1.205]; two-sided Wilcoxon signed-rank test, *W* = 0, *p <* .001). By treating the visual cue as substantially less reliable, forced fusion reduces its influence and can approximate the relatively flat, auditory-dominated responses produced by causal inference when precise sensory measurements favor separate causes. Thus, the reduced recovery does not indicate that the models make equivalent predictions under the same parameters; rather, forced fusion mimics causal-inference behavior through compensatory inflation of its visual-noise parameter. These simulations indicate that model identifiability is strongly constrained by sensory uncertainty and conflict magnitude. In particular, the empirical recovery analysis showed substantial confusion between forced fusion and causal inference, whereas probabilistic cue switching remained highly identifiable (84% correct recovery). Thus, within the experimentally tested conflict range and observed sensory-noise levels, competing integration models can generate highly similar behavioral predictions.

**Figure 21.**
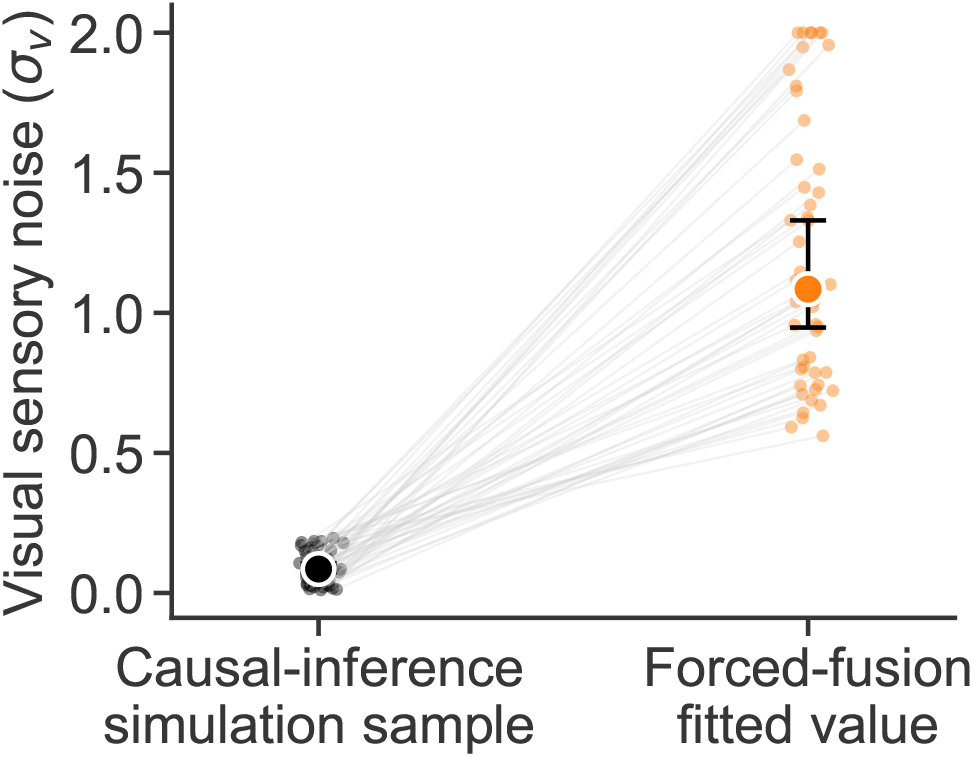
Visual-noise inflation allows forced fusion to mimic causal inference in the very-low-noise regime. Visual sensory noise used to generate datasets from the causal-inference model is compared with the value estimated when the same datasets were fitted by forced fusion. Results are shown for the wider maximum-conflict condition (0.45 s). Each connecting line represents one synthetic dataset (*N* = 50). Large circles indicate medians, and error bars show paired-bootstrap 95% confidence intervals for the medians. Forced fusion increased the fitted visual-noise parameter in every dataset, thereby reducing the effective contribution of the visual cue.

One implication of this analysis is that larger audiovisual conflicts could, in principle, improve model discriminability by producing stronger differences in likelihood and AIC across models. However, substantially increasing the conflict range was not feasible for the present experiment. First, audiovisual conflicts were defined relative to each participant’s subjective audiovisual bias, so that auditory and visual durations were adjusted to produce approximately matched perceived durations. For three participants we had a positive visual-duration bias, this calibration limited the range of negative conflicts that could be presented while keeping stimulus durations within the intended range. Second, extending the maximum conflict to approximately 0.45 s would have produced highly unusual audiovisual duration pairings. For example, at a +0.45 s conflict, the visual stimulus would approach 0.95 s, which may alter sensory uncertainty and produce behavior outside the regime targeted by the experiment. Thus, while larger conflicts may improve theoretical model recovery, they would also change the perceptual and experimental context being modeled.

